# Sotatercept Reverses SIN3a Deficiency-Driven PAH by Reprogramming BMPR2/TGF-β-HIF-1α Signaling Pathways

**DOI:** 10.64898/2026.02.03.703590

**Authors:** Katherine Jankowski, Anurupa Ghosh, Maria T. Ochoa, Shihong Zhang, Gregory David, Irene C. Turnbull, Malik Bisserier, Lahouaria Hadri

**Affiliations:** Department of Pharmacological Sciences, Icahn School of Medicine, Mount Sinai, New York, NY 10029, USA; Department of Cell and Molecular Physiology, New York Medical College, Valhalla, NY, 10595; Department of Cardiology, CVRI, Icahn School of Medicine, Mount Sinai, New York, NY 10029, USA; Department of Biochemistry and Molecular Pharmacology, NYU, NY 10016, USA

**Author notes:** Correspondence to Lahouaria Hadri, Department of Pharmacological Sciences, Icahn School of Medicine, Mount Sinai, New York, NY 10029, USA. Drs. Bisserier and Hadri are co-senior authors.

**Keywords:** Pulmonary arterial hypertension, BMPR2 signaling, TGF-β signaling, vascular remodeling, Sotatercept

## Abstract

**Background:** Pulmonary arterial hypertension is a progressive and fatal cardiopulmonary disease marked by excessive proliferation of pulmonary artery smooth muscle cells (PASMCs), pathological vascular remodeling, and ultimately right heart failure. Dysregulated BMPR2 signaling is a central molecular hallmark of PAH and is often associated with epigenetic suppression of BMPR2 expression. Switch-independent 3a (SIN3a), a transcriptional co-regulator and chromatin-modifying scaffold protein, has emerged as a key regulator of BMPR2 expression, yet its role in PAH pathogenesis remains poorly defined.

**Methods:** We generated smooth muscle cell-specific SIN3a knockout mice (SIN3a^SMC-/-^) and subjected them to the Sugen/hypoxia protocol to induce PAH. A cohort received Sotatercept treatment. In parallel, human PASMCs engineered to overexpress SIN3a were exposed to TGFβ1 or hypoxia (1% O₂) in vitro. Comprehensive transcriptomic profiling and pathway analyses identified molecular networks regulated by SIN3a and Sotatercept. Hemodynamic measurements and detailed morphometric analyses were used to assess disease severity and treatment response.

**Results:** SIN3a overexpression in PASMCs suppressed hypoxia-inducible factor-1α and TGF-β/SMAD2/3 signaling, restored BMPR2 expression, and activated canonical BMP signaling through SMAD1/5/9 phosphorylation, while reducing pro-inflammatory, oxidative, and fibrotic gene programs. Transcriptomic analyses revealed that SIN3a and Sotatercept converge on gene networks that regulate BMPR2 signaling, ID isoforms, extracellular matrix remodeling, oxidative stress, and inflammation. In vivo, smooth muscle–specific SIN3a deletion exacerbated Sugen/hypoxia-induced PAH, increasing right ventricular systolic pressure, right ventricular hypertrophy, pulmonary vascular remodeling, and fibrosis. Sotatercept treatment reversed these pathological features, restored SIN3a and BMPR2 expression, reactivated BMP signaling, and attenuated HIF-1α and TGF-β signaling in SIN3a-deficient mice.

**Conclusions:** SIN3a is a central epigenetic regulator of PASMC homeostasis that integrates oxidative stress, inflammation, and fibrotic signaling. Loss of SIN3a accelerates PAH progression, whereas Sotatercept restores SIN3a expression, rebalances BMPR2 and TGF-β signaling, and attenuates pulmonary vascular remodeling and right ventricular dysfunction. Together, these findings identify SIN3a as a disease-relevant therapeutic target and support the use of Sotatercept as a disease-modifying approach for pulmonary vascular disease.

## INTRODUCTION

Pulmonary arterial hypertension (PAH) is a progressive cardiopulmonary disease characterized by pulmonary vascular remodeling, increased pulmonary vascular resistance, and right ventricular failure. Despite advances in vasodilator-based therapies, PAH remains incurable, underscoring the need to identify disease-modifying mechanisms that directly target pathological vascular remodeling. A central molecular hallmark of PAH is the suppression of bone morphogenetic protein receptor type 2 (BMPR2) signaling. Reduced BMPR2 expression and activity are observed in both heritable and idiopathic PAH and contribute to excessive proliferation, resistance to apoptosis, and phenotypic dysregulation of pulmonary artery smooth muscle cells (PASMCs). In contrast, the activation of transforming growth factor-β (TGF-β) signaling promotes pro-fibrotic, inflammatory, and proliferative programs that further drive vascular remodeling. Disruption of the BMPR2/TGF-β signaling balance is a key determinant of disease progression. Emerging evidence indicates that BMPR2 suppression in PAH is not solely genetic but is strongly influenced by epigenetic mechanisms. PASMCs from patients with PAH exhibit stable, disease-associated transcriptional programs that persist ex vivo, suggesting durable epigenetic reprogramming. However, the specific chromatin regulatory mechanisms that integrate environmental stress signals, such as hypoxia and inflammation, with BMPR2 repression remain incompletely defined.

Switch-independent 3a (SIN3a) is a conserved epigenetic regulator and tumor suppressor that plays a critical role in normal development and cancer^1–7^. Notably, PAH shares several features with cancer, including hyperproliferation, resistance to apoptosis, and dysregulation of tumor suppressor pathways. SIN3a functions as a central scaffold of the SIN3 chromatin-modifying complex, coordinating transcriptional repression and activation in a context-dependent manner^8–10^. Because SIN3a lacks intrinsic DNA-binding activity, chromatin targeting depends on interactions with sequence-specific transcription factors and adaptor proteins^11–13^.

Our prior work identified reduced SIN3a expression in PASMCs from PAH patients, which was associated with BMPR2 promoter hypermethylation and transcriptional silencing. Restoration of SIN3a in vitro suppressed PASMC proliferation and enhanced BMPR2 expression and downstream BMP signaling. Moreover, lung-targeted delivery of SIN3a using aerosolized AAV1 improved hemodynamic parameters in experimental PAH models. Despite these observations, the mechanisms linking SIN3a to BMPR2 and TGF-β signaling crosstalk, particularly under hypoxic stress, remain unknown.

We hypothesized that SIN3a functions as a critical epigenetic regulator in PASMCs that maintains pulmonary vascular homeostasis by preserving BMPR2 signaling and limiting pathological activation of TGF-β and hypoxia-inducible factor 1 alpha (HIF-1α). To test this hypothesis, we generated a smooth muscle cell-specific SIN3a knockout mouse model and subjected it to the Sugen hypoxia model of PAH. Through integrated transcriptomic, molecular, and functional analyses, we demonstrated that the loss of SIN3a exacerbates PAH by destabilizing BMPR2 signaling, enhancing HIF-1α activity, and amplifying TGF-β-driven inflammatory, oxidative, and fibrotic responses. We further showed that treatment with Sotatercept attenuated pulmonary vascular remodeling and improved hemodynamic outcomes while restoring SIN3a expression, consistent with the suppression of HIF-1α–dependent pathogenic signaling. Together, these findings establish SIN3a as a central regulator of PASMC plasticity and identify epigenetic restoration of the SIN3a-BMPR2-TGF-β signaling axis as a disease-modifying mechanism in PAH, offering a compelling rationale for precise epigenetic therapies in PAH.

## MATERIAL AND METHODS

### Genetic Constructs and Pharmacological Treatments

SIN3a overexpression was achieved using an adenoviral vector encoding full-length human SIN3a (Ad-SIN3a) generated by VectorBuilder (Chicago, IL, USA). An adenovirus encoding green fluorescent protein (GFP) was used as a control (Ad-GFP). For pharmacological stimulation, recombinant human TGF-β1 (PeproTech) and Sotatercept (MedChem Express) were reconstituted according to the manufacturer’s instructions and used at final concentrations of 10 ng/mL and 5 nM, respectively, as per protocols established in previous studies.

### Cell Culture

Human pulmonary artery smooth muscle cells (PASMCs) from patients with PAH were obtained through the Pulmonary Hypertension Breakthrough Initiative (PHBI). Control PASMCs were isolated from failed-donor lungs deemed unsuitable for transplantation. Cells were cultured according to standard protocols and maintained in Smooth Muscle Growth Medium-2 (SmGM-2; Lonza) supplemented with 5% fetal bovine serum and smooth muscle cell growth supplements at 37°C in a humidified atmosphere containing 5% CO₂. PASMCs were used between passages 3 and 6 to ensure phenotypic consistency. All experiments were conducted on quiescent PASMCs that had been serum-starved in SmBM basal medium for 24 h prior to treatment. Hypoxic conditions were achieved by culturing cells at 1% O₂ with 5% CO₂, balanced with nitrogen, at 37°C in a humidified chamber. Cells were exposed to hypoxia for 24 hours unless otherwise specified.

### Western Blot

Total protein was extracted from PASMCs using radioimmunoprecipitation assay (RIPA) buffer (Thermo Fisher Scientific) supplemented with protease and phosphatase inhibitor cocktails (Thermo Fisher Scientific). Lysates were incubated on ice for 30 min with periodic vortexing and centrifuged at 15,000 × g for 20 min at 4°C to remove insoluble debris. Protein concentrations were determined using the bicinchoninic acid (BCA) assay (Thermo Fisher Scientific), and equal amounts of protein (50 µg) were run on 10-12% polyacrylamide gels. Proteins were transferred to nitrocellulose membranes using a semi-dry transfer system (Bio-Rad Trans-Blot Turbo) and blocked with 5% bovine serum albumin (BSA) or non-fat dry milk in Tris-buffered saline containing 0.1% Tween-20 (TBS-T) for 1 h at room temperature. The membranes were incubated overnight at 4°C with primary antibodies diluted in blocking buffer (antibody details are listed in **Supplementary Table 1**), then incubated for 1 h at room temperature with horseradish peroxidase (HRP)-conjugated secondary antibodies (Cell Signaling Technology). Immunoreactive bands were visualized using enhanced chemiluminescence (ECL; Thermo Fisher Scientific) and imaged using the ChemiDoc MP Imaging System (Bio-Rad). Densitometric quantification was performed using ImageJ software, with target protein levels normalized to β-actin or GAPDH, as appropriate.

### RNA Extraction and Quantitative Real-Time PCR

Total RNA was isolated from PASMCs using a RNeasy Mini Kit (Qiagen) according to the manufacturer’s protocol. RNA quantity and purity were assessed using a NanoDrop One spectrophotometer (Thermo Fisher Scientific), with A260/A280 ratios ranging from 1.9 to 2.1. cDNA synthesis was performed using qScript cDNA SuperMix (Quantabio). Quantitative PCR was conducted using PerfeCTa SYBR Green FastMix (Quantabio) on a QuantStudio 6 Pro Real-Time PCR system (Applied Biosystems), and each sample was analyzed in triplicate. Primer sequences targeting the human genes of interest were designed using Primer-BLAST; the sequences are listed in **Supplementary Table 1**. Relative transcript levels were calculated using the comparative threshold cycle (ΔΔCt) method and normalized to 18S rRNA or GAPDH, as reported in previous publications from our group. Data represent at least three independent biological replicates and are expressed as fold-change relative to control.

### Reactive oxygen species (ROS) measurement

Total intracellular reactive oxygen species were measured using the CellROX™ Orange Reagent (Invitrogen; Cat# C10443), a fluorescent probe for oxidative stress detection. PASMCs maintained under normoxic conditions, hypoxia (1% O₂), or TGF-β treatment, including SIN3a-overexpressing cells, were incubated with CellROX™ Orange Reagent at a final concentration of 5 µM in complete culture medium for 30 minutes at 37°C in the dark. Cells were washed with PBS and immediately analyzed by fluorescence microscopy. Upon oxidation by reactive oxygen species, CellROX™ Orange exhibits bright fluorescence with excitation and emission maxima at approximately 545 and 565 nm, respectively, enabling sensitive detection of intracellular oxidative stress.

### RNA Sequencing and Transcriptomic Analysis

Total RNA was isolated from PASMCs using an RNeasy Mini Kit (Qiagen) according to the manufacturer’s protocol. RNA integrity was assessed using an Agilent 2100 Bioanalyzer, and only samples with RNA integrity numbers (RIN) ≥7.0 and OD260/280 and OD260/230 ratios ≥2.0. RNA library preparation was performed using the TruSeq Stranded mRNA Library Prep Kit (Illumina), and sequencing was conducted on an Illumina NovaSeq 6000 platform (paired-end, 2 × 150 bp; Novogene, Sacramento, CA, USA). Each experimental condition was analyzed in triplicate. Raw sequencing reads were trimmed using Trimmomatic, aligned to the human genome (GRCh38) using the STAR aligner, and quantified at the gene level using featureCounts. Differential gene expression analysis was performed using the DESeq2 package. Genes were considered differentially expressed if they met the criteria of an adjusted p-value <0.05 (Benjamini-Hochberg FDR correction) and an absolute log₂ fold-change ≥1.5. Functional enrichment and regulatory inference were performed using multiple orthogonal approaches. Gene Ontology (GO) enrichment, KEGG pathway analysis, and transcription factor binding site prediction were performed using *ClusterProfiler* and *Enrichr*, enabling robust identification of perturbed biological processes and upstream regulatory networks. Transcriptomic data were uploaded for integrated Differential Expression and Pathway analysis (iDEP), which was used to perform regularized log transformation, principal component analysis (PCA), and unsupervised hierarchical clustering. Differentially expressed gene profiles were visualized using volcano plots, heatmaps, and PCA plots generated using iDEP. Z-score normalization of the top variable genes (n = 2,500) was used for heatmap generation. High-dimensional clustering and interactive pathway visualization were performed using *Clustergrammer* (http://amp.pharm.mssm.edu/clustergrammer/), following previously established methods. Transcript abundance was quantified using fragments per kilobase of transcript per million mapped reads (FPKM), which normalizes transcript length and sequencing depth. All RNA-seq datasets have been or will be deposited in a publicly accessible repository upon acceptance of this manuscript.

### Generation of tamoxifen-inducible Sin3a conditional knockout mice

Smooth muscle-specific, inducible deletion of Sin3a was achieved using a tamoxifen-inducible Cre-loxP system. Sin3a^flox/flox^ mice, in which exon 4 is flanked by loxP sites, were provided by Dr. Gregory David (New York University Langone Health). Excision of exon 4, which encodes a critical region of the SIN3a PAH2 domain, results in a frameshift and functional gene inactivation. To target SMC, Sin3a^flox/flox^ mice were crossed with SMMHC-CreERT2 transgenic mice (B6.FVB-Tg(Myh11-cre/ERT2)1Soff/J; Jackson Laboratory, #019079). Recombination was induced in adult male mice (8-10 weeks) by intraperitoneal tamoxifen administration (35 mg/kg/day in corn oil) for five consecutive days. Littermate controls lacked Cre or did not receive tamoxifen. Deletion efficiency was confirmed by quantitative PCR and Western blotting. All procedures were approved by the Institutional Animal Care and Use Committee of the Icahn School of Medicine at Mount Sinai. Deletion of Sin3a was confirmed prior to PAH induction by PCR-based genotyping, quantitative real-time PCR, Western blotting, and immunostaining for smooth muscle-specific assessment. Primer sequences used for genotyping and qPCR validation are listed in **Supplementary Table 1**.

### Sugen/Hypoxia-Induced PAH Model in Mice

PAH was induced in adult male C57BL/6J mice (8–10 weeks old; Jackson Laboratory) using the Sugen/hypoxia (SuHx) model, following protocols established in prior studies by Bisserier and Hadri. Mice were administered a once-weekly subcutaneous injection of SU5416 (20 mg/kg; MedChem Express), a vascular endothelial growth factor receptor 2 (VEGFR2) antagonist suspended in 0.5% carboxymethylcellulose. Immediately thereafter, the animals were housed in a normobaric hypoxia chamber (10% O₂) for 21 consecutive days using a digitally regulated ProOx 360 system (BioSpherix, Ltd.) with continuous oxygen monitoring and temperature control. Following hypoxia, the mice were returned to normoxia for an additional 7 days to allow disease stabilization. This model recapitulates key features of severe human PAH, including progressive vascular remodeling, right ventricular hypertrophy, and elevated right ventricular systolic pressure (RVSP). Age- and sex-matched control mice received vehicle injections and were maintained under normoxia. All animal procedures were conducted in accordance with the National Institutes of Health Guide for the Care and Use of Laboratory Animals and approved by the Institutional Animal Care and Use Committee at the Icahn School of Medicine at Mount Sinai.

### Sotatercept treatment of tamoxifen-Inducible SIN3a Conditional KO Mice

*SIN3a^SMC-/-^* KO and littermate mice were exposed to Sugen (SU5416) and hypoxic stress for 21 days. After 21 days, littermates and *SIN3a^SMC-/-^*KO mice received intraperitoneal (IP) injections of sotatercept at a dose of 10 mg/kg twice a week for 14 days. Hemodynamic measurements and tissue harvesting were performed on day 35.

### Right Ventricular Hemodynamic Assessment

RVSP was measured via open-chest catheterization, as previously described by Bisserier et al. ^8^. Briefly, mice were anesthetized with 2-4% isoflurane and maintained under 1-2% isoflurane using a precision vaporizer. The animals were positioned supine on a heated surgical platform to maintain body temperature at 37°C. Following tracheotomy, the mice were intubated with a 20G polyethylene catheter and mechanically ventilated (Harvard Apparatus MiniVent) at a tidal volume of 250 µL and a respiratory rate of 120 breaths/min. A midline sternotomy was performed to expose the heart. Using a dissecting microscope, the RV was visualized, and a 1.2F high-fidelity pressure catheter (Scisense, FTH-1212B-4518) was inserted directly through the RV free wall using a 30-gauge needle. The catheter was advanced into the right ventricle (RV) cavity, and real-time pressure tracings were recorded using a Scisense pressure catheter connected to an EMKA data-acquisition system. Hemodynamic signals were continuously monitored to confirm the stable and artifact-free RV waveforms. RVSP values were calculated as the peak systolic pressure over at least 10 cardiac cycles. At the end of the recording, the mice were euthanized by exsanguination under deep anesthesia, and tissue samples were collected for histological and molecular analyses. All analyses were performed in a blinded manner.

### Assessment of Right Ventricular Hypertrophy

Right ventricular hypertrophy was quantified using the Fulton index. Following completion of hemodynamic measurements, the hearts were excised and dissected into the RV and left ventricle plus septum (LV+S). Cardiac chambers were weighed, and the Fulton index was calculated as the RV/LV+S mass ratio. Tissues were subsequently processed for molecular and histological analyses.

### Wheat Germ Agglutinin (WGA) Staining

WGA staining was performed to delineate the cellular boundaries and quantify pulmonary vascular remodeling. Following euthanasia and transcardial perfusion with phosphate-buffered saline via the RV, RV tissue was excised, embedded in optimal cutting temperature (OCT) compound, and cryosectioned to a thickness of 5 μm. Tissue sections were incubated with Alexa Fluor 488–488-conjugated WGA (Thermo Fisher Scientific, Cat# W11261; 5 μg/mL in PBS) for 1 h at room temperature in the dark to label sialylated and N-acetylglucosaminylated glycoconjugates along the plasma membrane. Nuclei were counterstained with DAPI (1 μg/mL), and the sections were mounted using ProLong Gold Antifade Mountant (Thermo Fisher Scientific). Fluorescent images were acquired using an All-in-One Fluorescence Microscope BZ-X800 (Keyence) under standardized acquisition parameters across all experimental groups to ensure consistency of the results.

### Masson’s Trichrome Staining

RV cryosections (5 μm) were fixed in Bouin’s solution and stained using the Masson’s Trichrome Stain Kit (Sigma-Aldrich) to assess collagen deposition. Nuclei were stained with Weigert’s iron hematoxylin, muscle fibers with a Biebrich scarlet, and collagen with aniline. The sections were then dehydrated, mounted, and imaged under identical conditions. The fibrotic area (blue) was quantified using the ImageJ software and expressed as a percentage of the total RV area. Analyses were performed in a blinded manner.

### Vascular Morphometric Analysis

To assess pulmonary arterial remodeling, the lungs were inflated with a 1:1 mixture of PBS and optimal cutting temperature (OCT) compound, embedded in OCT, and snap frozen. Cryosections were cut at 8 μm thickness and fixed in cold acetone (−20°C) for 20 min. The sections were then rinsed in PBS and processed for hematoxylin and eosin (H&E) staining to visualize the vascular architecture. Morphometric analysis was performed on lung vessels from longitudinally sectioned distal arteries associated with the alveolar regions. Only small pulmonary arteries with external diameters ranging from 20 to 50 μm were included in the analysis. Digital images were captured under standardized magnification, and vessel measurements were performed in a blinded manner using ImageJ. The medial wall thickness was calculated using the following formula: % wall thickness = [(external diameter - internal diameter) / external diameter] × 100.

### Immunofluorescent Staining of Lung Sections

Immunofluorescent staining was performed to assess the localization and expression of target proteins in pulmonary tissues. Following euthanasia, the lungs were perfused with PBS via the RV, inflated with a 1:1 mixture of PBS and OCT compound, embedded in OCT, and snap-frozen. Cryosections were cut at 8 μm thickness and fixed in cold acetone (−20°C) for 20 min. The sections were permeabilized with 0.1% Triton X-100 in PBS for 10 min and blocked with 5% bovine serum albumin (BSA) in PBS for 1 h at room temperature. The primary antibodies diluted in the blocking solution were applied overnight at 4°C. After washing, the sections were incubated with the appropriate fluorophore-conjugated secondary antibodies for 1 h at room temperature in the dark. Nuclei were counterstained with DAPI (1 μg/mL), and the sections were mounted using ProLong Gold Antifade Mountant (Thermo Fisher Scientific). Fluorescent images were acquired using an All-in-One Fluorescence Microscope BZ-X800 (Keyence) under standardized acquisition parameters across all experimental groups. Image analysis was performed using ImageJ, and quantification was blinded. A complete list of the primary and secondary antibodies is provided in **Supplementary Table 1**.

### Statistical Analysis

Data are expressed as mean ± standard error of the mean (SEM). All statistical analyses were performed using GraphPad Prism (v10.0). Comparisons between two independent groups were evaluated using an unpaired two-tailed Student’s *t*-test. For experiments involving multiple conditions or variables, one-way or two-way ANOVA was applied, followed by Bonferroni’s multiple comparison test, where appropriate. Transcriptomic datasets were analyzed using the DESeq2 package, and significance thresholds were set at adjusted *p* < 0.05, using the Benjamini-Hochberg correction to control the false discovery rate. Statistical significance was set at p < 0.05.

## RESULTS

### SIN3a modulates global transcriptional programs in PASMCs

The development and progression of PAH are driven by a complex interplay between ECM remodeling, hypoxia-induced stress responses, and dysregulated TGF-β/BMPR2 signaling pathways, which collectively promote pathological pulmonary vascular remodeling ^8^. Emerging evidence indicates that epigenetic regulators play a central role in orchestrating the maladaptive transcriptional programs underlying these pathogenic processes in PAH ^14–17^.

To elucidate the transcriptional landscape regulated by SIN3a in PASMCs, we overexpressed SIN3a in human FD-PASMCs using an adenovirus vector and performed RNA sequencing followed by bioinformatic analyses. SIN3a overexpression resulted in a significant increase in SIN3a mRNA and protein levels compared with control cells (**Figure 1A**). Next, we performed RNA sequencing analysis in proliferating FD-PASMCs overexpressing SIN3a vs. controls (**Figure 1B**). Principal component analysis (PCA) revealed a clear separation between SIN3a-overexpressing and control PASMCs, indicating a substantial global transcriptomic reprogramming (**Supplementary Figure 1A**). Differential expression analysis demonstrated a widespread transcriptional reprogramming in response to SIN3a overexpression, as illustrated by the MA plot (**Supplementary Figure 1B**). Hierarchical clustering of the top 5,000 differentially expressed genes (DEGs) revealed distinct transcriptional modules corresponding to upregulated and downregulated gene sets (**Figure 1C**). Gene Ontology (GO) enrichment analysis identified four major biological clusters regulated by SIN3a (**Supplementary Figure 2A-B**). Cluster A comprised genes involved in cellular stress responses, including heat response and protein folding pathways. Cluster B was enriched for genes associated with RNA processing and ribonucleoprotein complex biogenesis. Cluster C included pathways related to G protein-coupled receptor signaling, ion transport pathways, and negative regulation of multicellular organismal processes. Notably, Cluster D was strongly enriched for extracellular matrix (ECM) remodeling and organization, cell adhesion, and biological adhesion pathways (**Supplementary Figure 2A-B**), highlighting a prominent role for SIN3a in ECM remodeling ^9^. Together, these data identify SIN3a as a key regulator of transcriptional networks governing stress response adaptation, RNA metabolism, and ECM architecture in PASMCs. Consistent with prior work demonstrating that SIN3a epigenetically regulates BMPR2 expression and PASMC proliferation, our findings further support a central role for SIN3a in pulmonary vascular remodeling and the molecular pathogenesis of PAH ^8,16,18^.

**Figure 1.**
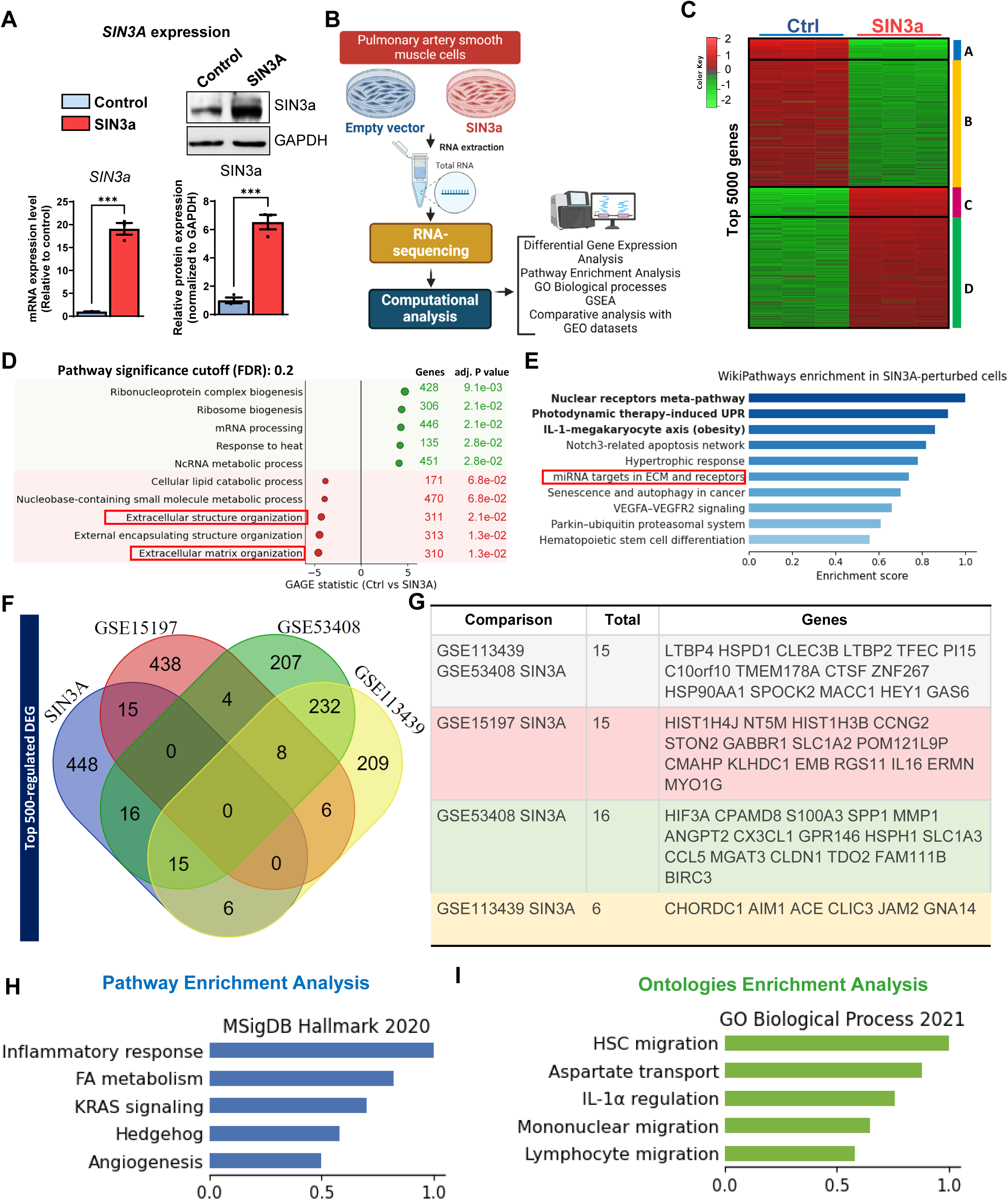
SIN3a overexpression induces global transcriptional programs in PASMCs. **(A)** Quantitative RT-PCR and immunoblot analysis confirming efficient SIN3a overexpression at the mRNA and protein levels in human proliferating failed donor FD-PASMCs following adenoviral transduction either with an empty vector (Ad-Control) or an adenoviral vector encoding human SIN3a (Ad-SIN3a) for 48 hours. **(B)** RNA sequencing workflow and analysis in SIN3a-overexpressing PASMCs and controls. (**C**) Heatmap of the top 5,000 DEGs illustrates a broad reprogramming of gene expression, consistent with SIN3a’s role as a master transcriptional regulator. **(D)** GAGE pathway enrichment analysis of differentially expressed genes reveals significant enrichment of pathways associated with TGF-β signaling, epithelial–mesenchymal transition (EMT), hypoxia, and ROS stress responses in SIN3a-overexpressing PASMCs. Functional interaction network of enriched pathways reveals SIN3a-driven upregulation (green) and downregulation (red) of signaling cascades, prominently involving extracellular matrix (ECM) dynamics and cellular stress responses**. (E)** WikiPathways enrichment identifies diverse regulatory modules following SIN3a overexpression, including nuclear receptor signaling and apoptotic pathways, implicating a multifaceted transcriptional role. (**F**) Cross-comparison with human PAH RNAseq datasets identifies a shared set of dysregulated genes modulated by SIN3a. Gene Set Enrichment Analysis (GSEA) demonstrates significant upregulation of gene sets linked to TGF-β signaling, epithelial-to-mesenchymal transition (EMT), and hypoxia. **(G-I)** Functional annotation of overlapping genes reveals enrichment in mitochondrial dysfunction, HIF1α activation, and TNF receptor signaling, implicating SIN3a in core disease-associated networks.

### SIN3a modulates epigenetically linked pathways and stress-adaptive signaling in PASMCs

SIN3a is a transcriptional scaffold protein that coordinates histone-modifying enzymes (HDACs) and repressor complex machinery, and plays a critical role in chromatin remodeling and gene regulation in both physiological and pathological contexts ^8,16,18,19^. Based on its established role in epigenetic regulation, we hypothesized that SIN3a gain-of-function exerts broad control over chromatin-associated transcriptional networks that contribute to PAH pathogenesis ^8,16,18^. To systematically test this hypothesis, we performed pathway-level enrichment analyses of DEGs identified in SIN3a-overexpressing FD-PASMCs. Using a stringent false discovery rate cutoff, GAGE (Generally Applicable Gene Set Enrichment) analysis revealed significant upregulation of canonical pathways implicated in pulmonary vascular pathology, including TGF-β signaling, epithelial-mesenchymal transition (EMT), hypoxia, and reactive oxygen species (ROS) stress response (**Figure 1D**), pathways previously shown to drive PASMC dysfunction and vascular remodeling in PAH ^20^. These findings are highly consistent with our prior transcriptomic and GO analyses and directly implicate SIN3a in regulating molecular programs governing cellular plasticity and oxidative injury^21^. Transcripts downregulated by SIN3a were significantly enriched for ECM-related processes, including extracellular structure organization and collagen biosynthesis, supporting the role of SIN3a in suppressing pro-fibrotic transcriptional programs associated with pulmonary vascular stiffening (**Figure 1D**). In contrast, upregulated genes were predominantly associated with mRNA processing and RNA splicing, implicating SIN3a in coordinating transcriptional and post-transcriptional regulation through chromatin-modifying complexes (**Figure 1D**) ^16^. To visualize the functional interplay among SIN3a-regulated pathways, we generated a network topology map depicting the convergence of upregulated (green) and downregulated (red) biological processes (**Supplementary Figure 3**). This network revealed extensive crosstalk among pathways involving ECM remodeling, transcriptional fidelity, and cellular stress responses, underscoring SIN3a’s role as an integrative regulator of vascular homeostasis. Further pathway interrogation using WikiPathways enrichment analysis identified SIN3a-regulated gene sets associated with nuclear receptor pathways, the unfolded protein response (UPR), apoptosis, and cardiac hypertrophy signatures (**Supplementary Figure 3**), all of which have been implicated in pulmonary vascular disease progression ^22–24^. Enrichment of autophagy, Parkin-ubiquitin-proteasome degradation, and miRNA-target regulatory modules further suggest that SIN3a influences PASMC behavior and growth through coordinated transcriptional and post-transcriptional mechanisms (**Figure 1E & Supplementary Figure 4)** ^25^. Collectively, these findings highlight SIN3a’s broad regulatory impact on PASMC transcriptional programs and stress-adaptive signaling pathways and support its central role in the molecular mechanisms driving PAH.

### SIN3a transcriptional targets converge on the pathogenic signaling modules implicated in PAH

The transcriptional landscape of PAH is regulated by multiple signaling events that drive vascular cell plasticity, inflammation, metabolic dysregulation, and ECM remodeling ^26^. To identify gene networks specifically modulated by SIN3a in this context, we performed refined gene set enrichment analyses on the top 250 upregulated and downregulated transcripts in proliferating SIN3a-overexpressing FD-PASMCs. Using the Hallmark Molecular Signatures and GO databases, we found that SIN3a-upregulated genes were significantly enriched in pathways associated with stress-adaptive and vascular remodeling (**Supplementary Figure 4A-5A**). These include EMT, hypoxia signaling, glycolysis, cholesterol biosynthesis, and apoptosis (**Supplementary Figure 4A-5A**), all of which have been implicated in PASMC phenotypic switching and pulmonary vascular remodeling in PAH ^27^. These pathways are well-established drivers of PASMCs’ hyperproliferative, pro-inflammatory activation, and vascular remodeling in human PAH ^27^. In contrast, the top 250 downregulated genes in SIN3a-overexpressing PASMCs were enriched in pathway signatures of unresolved cellular stress, including the unfolded protein response (UPR) signaling, IL2/IL6 cytokine signaling, and chronic hypoxic injury response (**Supplementary Figure 4B-5B**), suggesting attenuation of maladaptive stress. In addition, SIN3a-modulating transcripts showed significant enrichment in pro-inflammatory signaling cascades, particularly the IL6-JAK-STAT3 and TNF-α/NF-κB pathway axes (**Supplementary Figure 6).** These findings underscore the context-dependent regulatory role of SIN3a, whereby it dampens excessive inflammatory and stress-associated transcriptional programs while promoting adaptive or compensatory signaling responses under cellular challenge.

To assess the clinical relevance of SIN3a-regulated gene networks, we performed an integrative transcriptomic comparison between our *in vitro* PASMC dataset and publicly available RNA-seq profiles from human PAH lung tissue (**Figure 1F**). This analysis identified a subset of overlapping transcripts that were both dysregulated in human PAH lung tissue and modulated by SIN3a in PASMCs, indicating a functional convergence on disease-relevant molecular signatures. Subsequent pathway and ontology analyses of these shared transcripts revealed significant enrichment in inflammatory signaling, mitochondrial dysfunction, and apoptotic processes (**Figure 1G**). Notably, key pathways included the SOD/TNFR1 signaling axis, hypoxia-inducible factor (HIF1α) signaling, and TNF receptor-mediated inflammatory responses, all of which are mechanistically linked to pulmonary vascular remodeling and PAH pathogenesis (**Figure 1G-I**). Collectively, these findings position SIN3a as an upstream regulator of transcriptional networks that intersect both canonical and non-canonical pathways involved in PAH, thereby providing a mechanistic link between epigenetic regulation at the chromatin level and the molecular drivers of pulmonary vascular disease.

### SIN3a represses HIF1α signaling and restores the TGF-β/BMPR2 axis in PASMCs under hypoxic stress

In PAH, the activation of HIF1α and TGF-β signaling promotes PASMC proliferation, ECM remodeling, and inflammation ^28,29^. HIF1α, which is stabilized under low hypoxic conditions, transcriptionally regulates fibrotic and glycolytic programs, whereas TGF-β signaling suppresses BMPR2 signaling, a hallmark feature of heritable and idiopathic PAH ^30^. Based on our transcriptomic data, we hypothesized that SIN3a functions as a negative regulator of the HIF1α/TGF-β/BMPR2 signaling axis.

To test this hypothesis, primary human failed donor (FD)-PASMCs were cultured under hypoxic conditions (1% O₂) and treated with recombinant TGF-β1, in the presence or absence of SIN3a overexpression, followed by the assessment of gene expression. Notably, exposure to hypoxia and TGF-β1 treatment significantly reduced *SIN3a* mRNA levels (**Figure 2A**), consistent with our previous observations in PAH patients and the SuHx mouse model ^8,16^. Together, these findings indicate a disease-relevant downregulation of SIN3a in response to fibrotic and hypoxic stress, supporting its role as a key regulator of PAH pathogenesis. Similarly, TGF-β1 expression was potentiated by both hypoxia and TGF-β1 stimulation, indicative of positive feedback that amplifies pathological signaling (**Figure 2B**). Notably, SIN3a overexpression significantly suppressed *TGF-β1* levels, suggesting a repressive role of this feedback mechanism (**Figure 2B**). In contrast, *BMPR2*, a protective signaling component frequently downregulated in PAH, was suppressed by both hypoxia and TGF-β1, but its expression was restored by SIN3a overexpression (**Figure 2C**). To examine whether SIN3a modulates transcriptional responses to hypoxia and TGF-β, we analyzed publicly available chromatin immunoprecipitation followed by sequencing (ChIP-seq) datasets from ENCODE and ReMap. These analyses identified prominent HIF1α binding peaks in the promoter and intragenic regions of *TGFB1*, *TGFBR1*, *SMAD3*, and *SMAD4* (**Figure 2D** & **Supplementary Figure 7**), suggesting the transcriptional regulation of TGF-β signaling components by hypoxia-responsive elements. As expected, *HIF1A* transcript levels were induced by hypoxia and further enhanced by TGF-β1 treatment (**Figure 2E**). Notably, SIN3a overexpression markedly reduced *HIF1A* mRNA expression levels induced by *TGFB1* and under all conditions, indicating the transcriptional repression of this key hypoxia effector by SIN3a (**Figure 2E**). Immunoblot analysis of whole-cell lysates confirmed these transcriptional changes at the protein level. Hypoxic conditions and TGF-β1-treated PASMCs exhibited increased HIF1α and reduced BMPR2 (**Figure 2F**). In contrast, in hypoxic conditions, SIN3a overexpression restored BMPR2 protein levels while reducing HIF1α protein levels in cells treated with TGF-β1 (**Figure 2F**). Next, we assessed the subcellular localization of HIF1α in proliferating FD-PASMCs by immunofluorescence. Under hypoxic conditions and TGF-β1 stimulation, we observed that SIN3a overexpression reversed the nuclear translocation of HIF1α, a hallmark of its transcriptional activation, as well as SMAD2/3 nuclear phosphorylation, key mediators of TGF-β signaling (**Figure 2G**), highlighting its ability to simultaneously suppress pathogenic TGF-β pathway activity while promoting BMPR2-mediated signaling (**Figure 2F-G**). Finally, to evaluate the downstream functional impact of SIN3a-mediated transcriptional modulation, we quantified the expression of canonical markers of oxidative stress, ECM remodeling, and inflammation by qRT-PCR. Specifically, *NOX4* and *NFE2L2* were selected as representatives of ROS-generating and antioxidant pathways, respectively; *COL1A1* and *COL3A1* reflect collagen-driven matrix remodeling; and IL-6 captures pro-inflammatory cytokine activation. All five transcripts were markedly downregulated in SIN3a-overexpressing cells (**Figure 2H-I & Supplementary Figure 8**), demonstrating the ability of SIN3a to suppress hypoxia- and TGF-β-induced pathogenic transcriptional programs. Furthermore, to confirm that SIN3a attenuates hypoxia (0.1% O₂)-induced oxidative stress, we performed a complementary assay to measure the total cellular ROS. As expected, ROS levels were significantly elevated under hypoxic and TGF-β conditions compared to normoxia, and these effects were reversed by SIN3a overexpression (**Figure 2J**). Together, these findings established that SIN3a is a critical regulator of PASMC stress adaptation in vitro. By suppressing HIF1α transcription and activity, repressing TGF-β signaling, and restoring BMPR2 expression and SMAD1/5/9 signaling, SIN3a attenuates multiple intersecting molecular pathways that drive pulmonary vascular remodeling in PAH.

**Figure 2.**
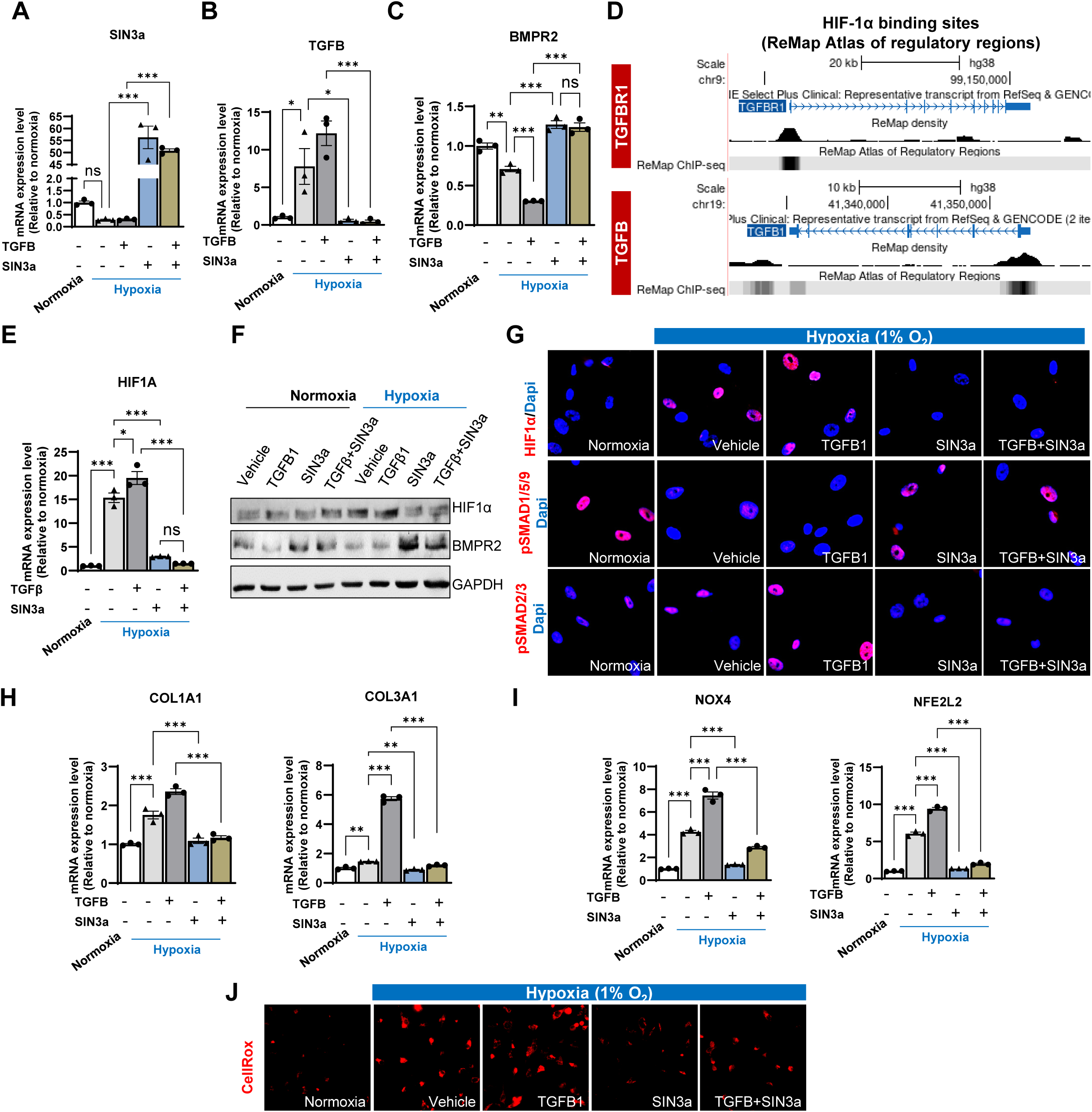
SIN3a modulates hypoxia- and TGF-β-induced signaling pathways in PASMCs. **(A)** (**A**) qRT-PCR analysis of SIN3a mRNA expression in PASMCs cultured under normoxic conditions or exposed to hypoxia (1% O₂) and/or TGF-β1 stimulation. (**B-C**) qRT-PCR analysis of TGFB1 and BMPR2 mRNA expression under hypoxic and/or TGF-β1 conditions in the presence or absence of SIN3a overexpression. (**D**) ENCODE ChIP-seq analysis demonstrating HIF-1α binding at promoter regions of TGF-β signaling components, including TGFBR1 and TGFB1. (**E**) qRT-PCR analysis of HIF-1α mRNA expression under hypoxia and/or TGF-β1 stimulation, with or without SIN3a overexpression. (**F**) Representative immunoblot analysis of HIF-1α and BMPR2 protein expression under identical experimental conditions. (**G**) Immunofluorescence staining showing nuclear localization of HIF-1α and phosphorylation of SMAD1/5/9 under hypoxic and TGF-β1 conditions, demonstrating attenuation of hypoxia-induced nuclear translocation by SIN3a. (**H-I**) qRT-PCR analysis of COL1A1, COL3A1, NOX4, and NFE2L2 mRNA expression in PASMCs exposed to hypoxia (1% O₂) and TGF-β1, with or without SIN3a overexpression. (**J**) Reactive oxygen species detection in PASMCs cultured under normoxic or hypoxic (1% O₂) conditions using CellROX reagent (5 μM) added during the final 30 min of treatment. Cells were fixed, and CellROX fluorescence intensity (red) was quantified as a measure of intracellular oxidative stress. Data are presented as mean ± SEM. P < 0.05, P < 0.01, P < 0.001.

### Smooth muscle cell-specific deletion of SIN3a exacerbates PH and right ventricular remodeling in vivo

To further investigate the functional significance of SIN3a *in vivo* within a lineage-restricted context, we generated a tamoxifen-inducible SMC-specific *SIN3a* knockout mouse model (*Sin3a^fl/fl^; SMMHC-^CreER^T^*^2^, hereafter referred to as *SIN3a^SMC-/-^*). Efficient and cell type-specific deletion of SIN3a in VSMC was confirmed via genomic PCR, qRT-PCR, and immunostaining analyses of SIN3a in aortic tissue (**Supplementary Figure 9A & Figure 3A-B**). Importantly, the expression of *SIN3b* isoform remained unchanged (**Supplementary Figure 9B),** establishing a robust model for assessing the consequences of SIN3a deletion in the pulmonary vasculature. *SIN3a^SMC-/-^* and littermate *SIN3a^fl/fl^*controls were subjected to the SuHx protocol, a well-established protocol that mimics the key histopathological features of human PAH, including occlusive vascular remodeling and RV hypertrophy ^31^. Notably, right heart catheterization revealed significantly elevated RVSP in *SIN3a^SMC-/-^* mice under normoxic conditions, suggesting the development of spontaneous PAH features (**Figure 3C, left panel**). Following three weeks of SuHx exposure, RVSP was further increased in *SIN3a^SMC-/-^* KO mice compared with littermate controls (**Figure 3C, left panel).** Consistent with these hemodynamic changes, *SIN3a^SMC-/-^*mice subjected to SuHx developed a marked RV hypertrophy as evidenced by an increased Fulton index (RV/[LV+S]) (**Figure 3C, right panel**). To further characterize the extent of cardiac remodeling, we assessed RV cardiomyocyte hypertrophy by quantifying cardiomyocyte cross-sectional area. SMC-specific *SIN3a* deletion resulted in significantly enlarged RV cardiomyocytes under SuHx conditions compared with normoxic controls (**Figure 3D**). This hypertrophic response was accompanied by marked upregulation of hypertrophy canonical hypertrophy-associated genes, including *ANP*, *BNP*, and *β-MHC*, in the RV tissue, as measured by qRT-PCR (**Figure 3E**). These markers are well-established indicators of maladaptive ventricular remodeling in both clinical and preclinical studies ^32^. Masson’s trichrome staining revealed increased collagen deposition in the RV sections of *SIN3a^SMC-/-^* mice exposed to SuHx (**Figure 3F**). These histopathological analyses corroborated with the transcriptomic changes in the levels of pro-fibrotic markers such as *TGFβ1*, *COL1A1*, and *COL3A1*in the RV of *SIN3a^SMC-/-^* mice following SuHx exposure (**Figure 3G**), implicating SIN3a in the regulation of fibrotic gene programs and supporting the presence of interstitial fibrosis as a key component of RV remodeling in this model. Our *in vivo* data provided compelling evidence that specific SMC SIN3a deficiency exacerbates the severity of experimental PH, highlighting the importance of epigenetic regulation in PAH pathogenesis.

**Figure 3.**
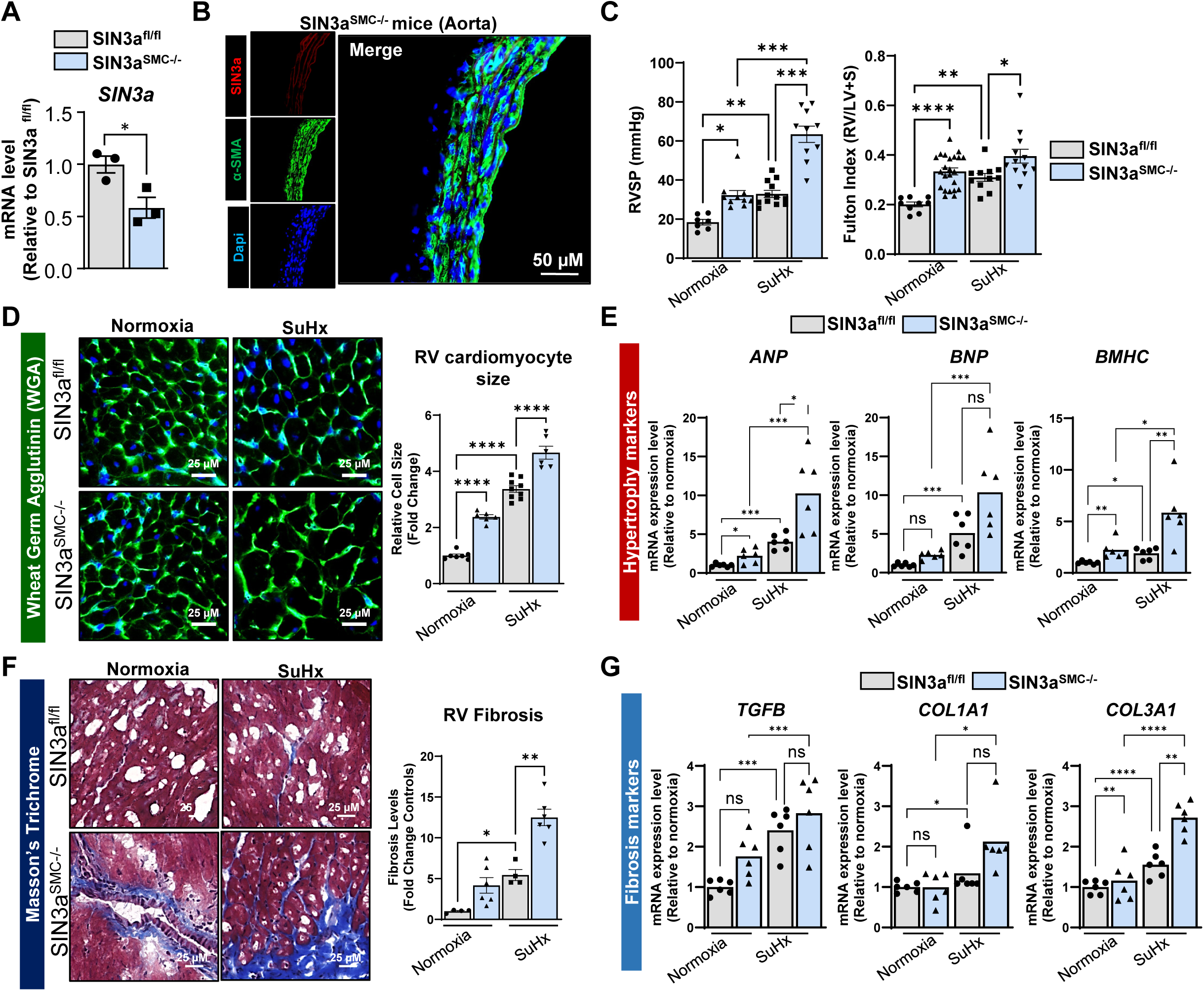
Smooth muscle-specific SIN3a deficiency exacerbates RV function and remodeling in SuHx-induced PH. **(A)** SIN3a mRNA expression in aortic tissue from tamoxifen-induced SMMHC-CreERT2 SIN3a^fl/fl^ mice (SIN3a*^SMC-/-^*) mice and littermates SIN3a*^fl/fl^*control mice. **(B)** Representative co-immunofluorescence staining of SIN3a (red) and α-smooth muscle actin (α-SMA; green) in aortic sections of SIN3a*^SMC-/-^* mice. (**C)** Right heart catheterization analysis showing right ventricular systolic pressure (RVSP; left panel) in SIN3a*^SMC-/-^* KO and littermate control mice under normoxic and Sugen/Hypoxia (SuHx) conditions for 21 days. RV hypertrophy was quantified using the Fulton index (RV/LV+S), right panel. **(D)** Quantification of cardiomyocyte cross area assessed by wheat germ agglutinin (WGA) staining in SIN3a*^SMC-/-^* KO and littermate control mice under normoxic and SuHx conditions. **(E)** qRT-PCR analysis of pro-hypertrophic gene markers (*ANP, BNP, BMHC*) in RV from SIN3a-deficient mice compared to littermates under normoxic and SuHx conditions. **(F)** Representative Masson’s trichrome staining of RV sections of SIN3a*^SMC-/-^* KO mice and littermate control mice under normoxic and SuHx conditions. **(G)** Fibrosis-related gene expression (*TGFB, COL1A1, COL3A1*) was measured by qRT-PCR in RV tissues from SIN3a*^SMC-/-^* KO mice compared to littermates exposed to normoxic and SuHx conditions. Data are expressed as mean ± SEM. *p < 0.05, **p < 0.01, ***p < 0.001.

### Loss of SIN3a in smooth muscle cells promotes pulmonary vascular remodeling, inflammation, and oxidative stress in the lungs of SuHx-induced PH mice

We next confirmed the efficiency and specificity of SIN3a deletion in PASMCs using immunofluorescence staining for SIN3a and α-smooth muscle actin (α-SMA) in lung sections from *SIN3a^fl/fl^* and *SIN3a^SMC–/–^* mice under normoxic or SuHx conditions. SIN3a expression was significantly reduced under SuHx conditions in both *SIN3a^fl/fl^* and *SIN3a^SMC–/–^* mice. Importantly, Sin3a was nearly absent in muscularized α-SMA-positive cells in *SIN3a^SMC–/–^* mice, confirming the SMC-specific depletion of SIN3a (**Figure 4A**). To determine whether SIN3a loss modulates vascular remodeling, we performed histomorphometric analysis of the distal pulmonary arteries in *SIN3a^SMC-/-^* and littermate controls and found that *SIN3a^SMC-/-^* mice displayed a marked increase in medial wall thickness compared to controls under SuHx, consistent with the exacerbated muscularization of distal arterioles (**Figure 4B**). Notably, these pathological changes closely mirrored those observed in the lungs of patients with PAH, underscoring the functional role of SIN3a in restraining the structural remodeling of the pulmonary vasculature. Through epigenetic and transcriptional mechanisms, SIN3a may maintain chromatin homeostasis and repress pro-inflammatory mediators and pro-fibrotic gene programs. Indeed, the loss of SIN3a disrupts this balance, leading to elevated levels of inflammatory cytokines, such as TNF-α, which amplify the signaling cascades that promote fibroblast activation (**Figure 4D & Supplementary Figure 10**). Concurrently, the expression of *TGFB1, COL1A1,* and *COL3A1 was* increased (**Figure 4C**), which drives ECM deposition and tissue stiffening. The upregulation of α-SMA and SNAI1 transcripts without a change in SLUG (**Supplementary Figure 11**) further indicates enhanced myofibroblast differentiation and/or EMT transition, processes central to pathological remodeling in PAH ^33^. Collectively, these findings underscore SIN3a’s function as a key epigenetic regulator whose deficiency exacerbates inflammatory and fibrotic burdens. Next, we evaluated redox-associated gene expression to determine whether SIN3a regulated oxidative homeostasis *in vivo*. Loss of SIN3a led to significant upregulation of *NOX4*, a pro-oxidant NADPH oxidase enzyme that promotes ROS generation, accompanied by downregulation of the antioxidant defense genes *SOD2* and *NFE2L2* (**Figure 4E**). These alterations indicate that SIN3a helps to maintain oxidative homeostasis in the pulmonary vasculature, a process tightly linked to vascular remodeling and endothelial dysfunction in PAH ^34^. Consistent with these findings, expression analysis of PAH-associated signaling pathways revealed significantly reduced *BMPR2* mRNA levels and increased *HIF1A* transcript levels in *SIN3a^SMC-/-^* lungs, confirming our *in vitro* observations (**Figures 4F-G**). Importantly, these transcriptional changes were corroborated at the protein level: SIN3a deletion reduced BMPR2 protein expression and diminished SMAD1/5/9 phosphorylation, indicative of impaired canonical BMP signaling. In contrast, the protein levels of HIF1α and phospho-SMAD2/3 were elevated, reflecting enhanced hypoxia and TGF-β signaling (**Figure 4H-J**). Together, these findings provide compelling *in vivo* evidence that SIN3a is a central regulator of pulmonary vascular homeostasis by coordinating anti-inflammatory, anti-fibrotic, antioxidant, and BMP-promoting transcriptional programs.

**Figure 4.**
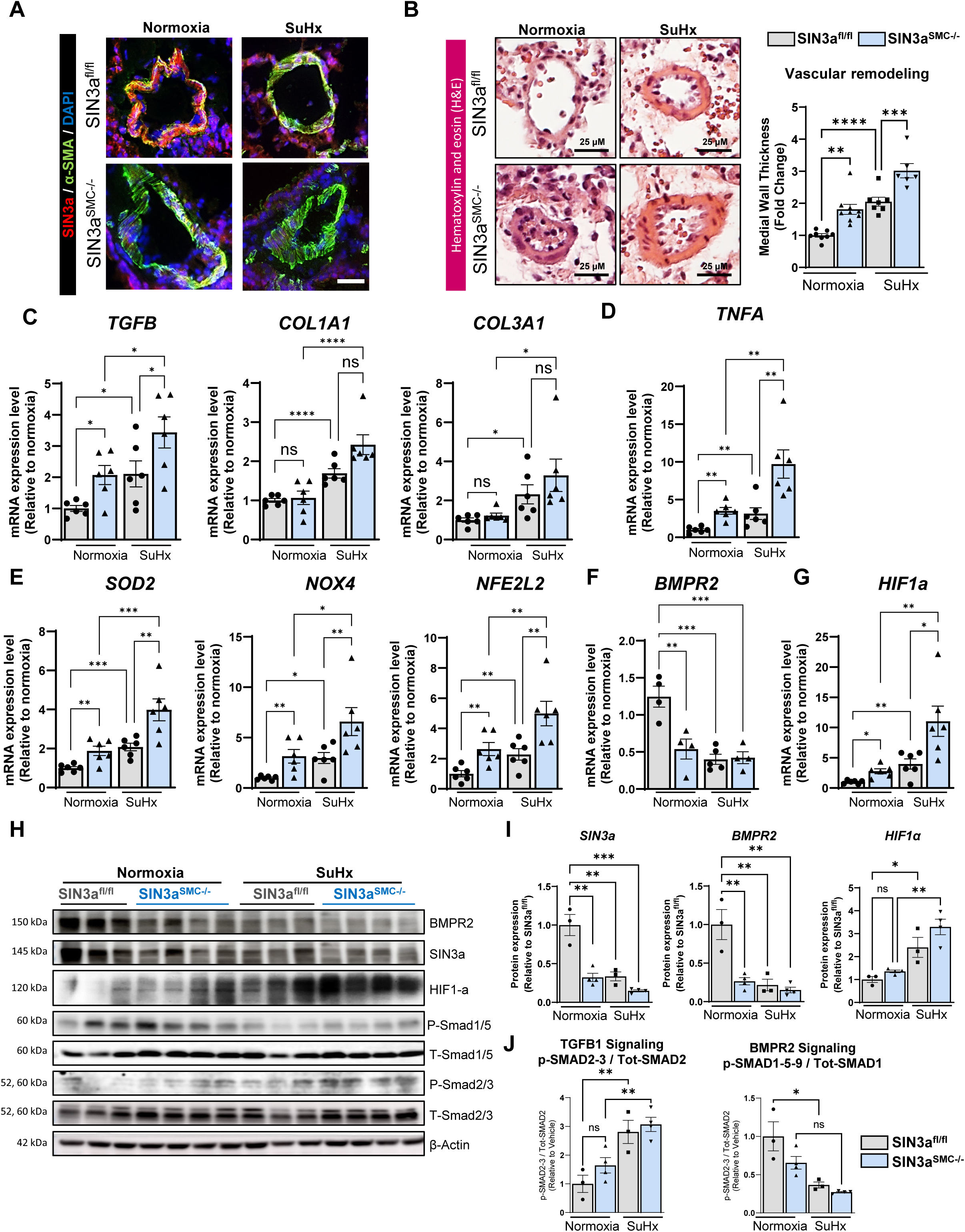
Loss of SIN3a promotes inflammation, oxidative stress, and vascular remodeling in SuHx-induced PH. **(A)** Representative co-immunofluorescence staining of α-smooth muscle actin (α-SMA; green) and SIN3a (red) in lung sections from littermate control and SIN3a*^SMC-/-^*mice under normoxic and SuHx conditions. **(B)** Quantification of pulmonary arterial medial wall thickness and vascular remodeling in SIN3a*^SMC-/-^*mice littermate control and SIN3a*^SMC-/-^* mice under normoxic and SuHx conditions, as assessed by hematoxylin and eosin staining. **(C-D)** qRT-PCR analysis of inflammatory (*TNFA*) and profibrotic markers (TGFB, COL1A1, and COL3A1) in lungs from littermates and SIN3a*^SMC-/-^* mice under to normoxic and SuHx conditions. **(E)** qRT-PCR analysis of oxidative stress–related genes (*SOD2, NOX4, NFE2L2*) in lungs from controls littermates and SIN3a*^SMC-/-^* mice under normoxic and SuHx conditions. **(F-G)** qRT-PCR analysis of *BMPR2* and *HIF1A* transcript levels in lungs of littermates and SIN3a*^SMC-/-^* mice under normoxic and SuHx conditions. **(H)** Representative immunoblot analysis of BMPR2, phospho-SMAD2/3, and phospho-SMAD1/5/9 protein levels in lungs from SIN3a*^SMC-/-^*lungs. **(I-J)** Densitometric quantification of immunoblot data showing SIN3a and BMPR2 protein levels normalized to β-actin, and phospho-SMAD2/3 (TGF-β signaling) and phospho-SMAD1/5/9 (BMP signaling) normalized to total SMAD2/3 and SMAD1, respectively. Data are presented as mean ± SEM. *p < 0.05, **p < 0.01, ***p < 0.001, ns: not significant.

### Global transcriptomic changes induced by SIN3a overexpression and SIN3a together and in combination with Sotatercept in PAH-PASMCs

Sotatercept is a first-in-class fusion protein comprised of the extracellular domain of human activin receptor type IIA (ActRIIA) linked to the Fc domain of human IgG1. It functions as a ligand trap for TGF-β superfamily members, most notably activins and growth differentiation factors (GDFs), effectively reestablishing the balance between the proliferative activin/GDF pathway and the anti-proliferative BMP pathway ^35,36^. In PAH, disruption of this signaling equilibrium promotes the excessive proliferation of PASMCs. By sequestering circulating activins, sotatercept enhances BMP signaling through BMPR2, redirecting cellular responses toward growth control and attenuating disease progression ^37,38^.

Given that SIN3a plays an independent role in modulating the TGF-β/BMPR2 axis, we investigated its effects alone and in combination with Sotatercept on gene expression in PAH-PASMCs using RNAseq to comprehensively assess treatment-induced transcriptomic changes. PCA revealed clear separation between SIN3a-overexpressing and control PAH-PASMCs, with partial overlap observed in cells treated with Sotatercept, indicating substantial global transcriptomic remodeling in response to SIN3a and Sotatercept (**Figure 5A**). Heatmap visualization and differential expression analysis further identified a distinct transcriptional signature compared with the controls (**Figure 5B**). Unsupervised hierarchical clustering of significantly differentially expressed genes (DEGs) demonstrated robust segregation between SIN3a- and SIN3a + Sotatercept-treated cells and control PAH-PASMCs (**Figure 5C**). Functional analysis revealed coordinated regulation of genes involved in proliferative signaling, with upregulation of SIN3a and genes associated with cellular homeostasis, hypoxic adaptation, and stress responses. In parallel, genes linked to pro-fibrotic TGF-β signaling were consistently downregulated following SIN3a overexpression and combined SIN3a + Sotatercept treatment (**Figure 5C-D**). Collectively, GO enrichment analyses supported a multifaceted role for Sotatercept in modulating pathogenic signaling, stress adaptation, and structural remodeling in PAH-PASMCs.

**Figure 5.**
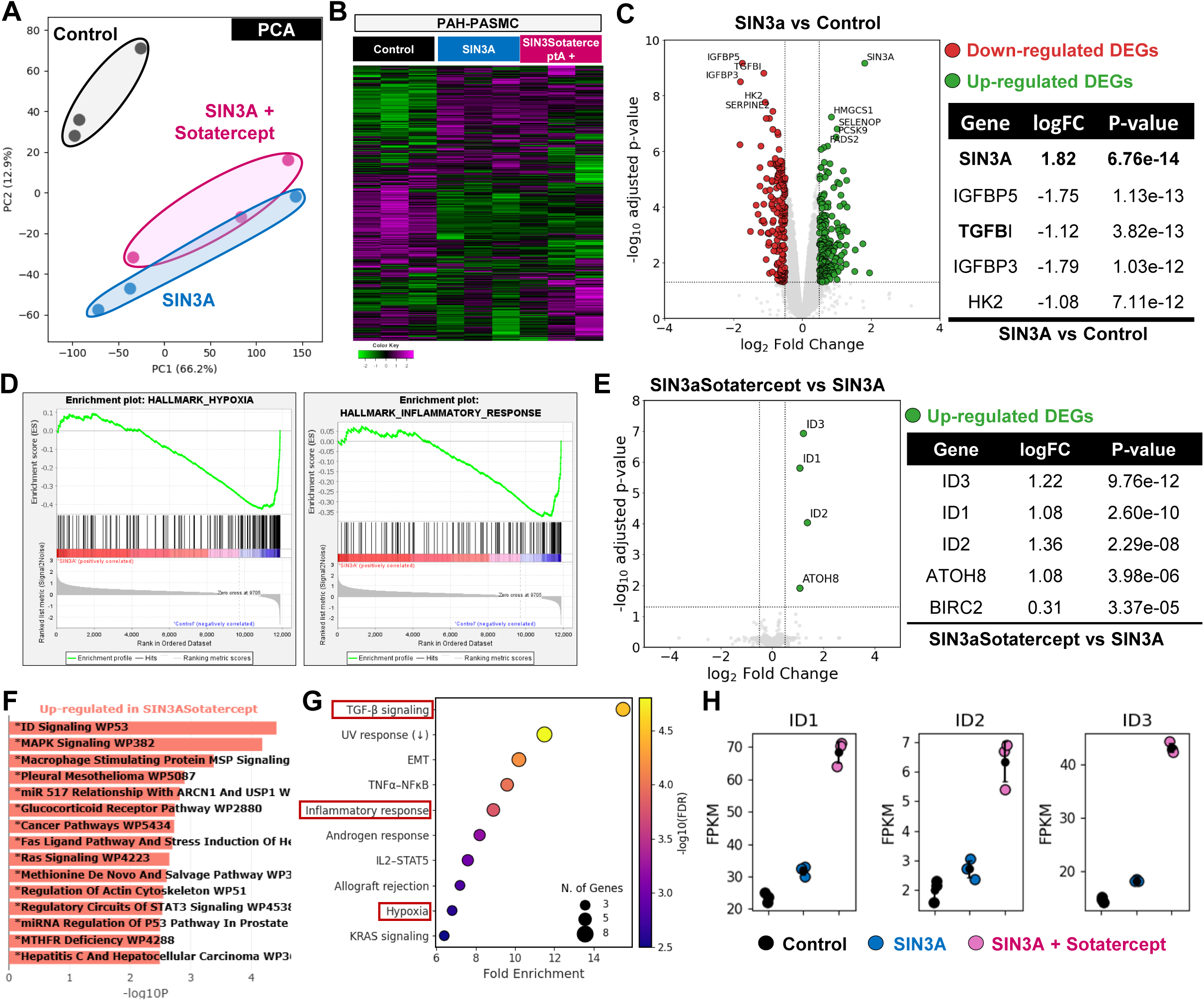
**Global transcriptomic profiling of PAH-PASMCs following SIN3a overexpression and combined SIN3a with Sotatercept treatment**. (**A**) Principal component analysis (PCA) of RNA-seq data showing distinct clustering of control, SIN3a-overexpressing, and SIN3a + sotatercept-treated PAH-PASMCs. (**B**) Heatmap of significantly differentially expressed genes (DEGs) across experimental conditions. (**C**) Unsupervised hierarchical clustering illustrating treatment-specific transcriptional signatures and regulation of genes associated with proliferative signaling, cellular homeostasis, stress response, and TGF-β signaling. (**D**) Gene set enrichment analysis (GSEA) reveals enrichment for hypoxia- and oxidative stress-related pathways. (**E**) Differential expression of BMPR2 downstream target genes ID1, ID2, and ID3 following SIN3a and SIN3a + Sotatercept treatment. (**F-G**) Gene Ontology (GO) enrichment analyses highlighting biological processes related to vascular remodeling, ECM organization, hypoxia response, oxidative stress, and PASMC proliferation. (**H**) FPKM-normalized expression levels of ID1, ID2, and ID3 across control, SIN3a-overexpressing, and SIN3a + Sotatercept-treated PAH-PASMCs.

Among the top DEG, SIN3a and Sotatercept jointly regulated pathways involving insulin-like growth factor-binding proteins, including IGFBP3 and IGFBP5, which are key modulators of insulin-like growth factor (IGF) signaling and influence cell growth, differentiation, survival, and fibrotic responses (**Figure 5C**) ^39,40^. These findings indicate that SIN3a and Sotatercept induce a reproducible, treatment-specific transcriptional program in PAH-PASMCs consistent with the attenuation of pathological vascular remodeling and fibrosis. GSEA demonstrated a significant enrichment of pathways related to hypoxia signaling and oxidative stress responses (**Figure 5D**). Notably, combined SIN3a and Sotatercept treatment was associated with increased expression of ID1, ID2, and ID3, canonical downstream targets of BMPR2 signaling that exert anti-proliferative effects (**Figure 5E**). GO analysis of DEGs further highlighted significant enrichment in biological processes related to vascular remodeling, including TGF-β receptor signaling, ECM organization, hypoxia response, oxidative stress response, and SMC proliferation (**Figure 5F-G**). Comparative transcriptomic analysis revealed differential expression of BMP target genes ID1, ID2, and ID3 across control, SIN3a-overexpressing, and SIN3a plus Sotatercept-treated PAH-PASMCs, consistent with the modulation of canonical BMP signaling (**Figure 5H**). In summary, RNA-sequencing analyses demonstrated that SIN3a overexpression alone and in combination with Sotatercept profoundly reshaped the transcriptomic landscape of PAH-PASMCs by suppressing TGF-β-driven ECM remodeling, modulating hypoxia and oxidative stress pathways, and restoring gene programs associated with vascular homeostasis. These findings provide mechanistic insights into the therapeutic effects of Sotatercept in PAH.

### Sotatercept treatment attenuates PAH features in SIN3a*^SMC-/-^* KO mice by restoring the BMPR2/TGFβ balance and reducing HIF1α expression

Since Sotatercept acts as a ligand trap to rebalance BMPR2/TGFβ signaling pathways, we next evaluated its therapeutic efficacy in SIN3a*^SMC-/-^*KO *in vivo* using the SuHx PAH model. Starting on day 21, mice received intraperitoneal injections of Sotatercept (10 mg/kg) or vehicle twice weekly for 14 days (**Figure 6A**). Remarkably, Sotatercept administration significantly attenuated maladaptive RV remodeling as evidenced by reduced RVSP and RV hypertrophy (lower Fulton index) compared to vehicle (**Figure 6B**). Furthermore, Sotatercept markedly diminished RV cardiomyocyte hypertrophy, the expression of cardiac hypertrophy-associated markers, RV fibrosis, and profibrotic markers in SIN3a^SMC-/-^ KO mice compared with vehicle-treated mice (**Figure 6C-F**). Histological analysis revealed a significant reduction in distal pulmonary vascular remodeling (**Figure 6G**). In addition, Sotatercept reduced profibrotic, inflammatory, and oxidative markers in the lungs of SIN3a*^SMC-/-^*KO mice compared to controls (**Figure 6H-J**).

**Figure 6.**
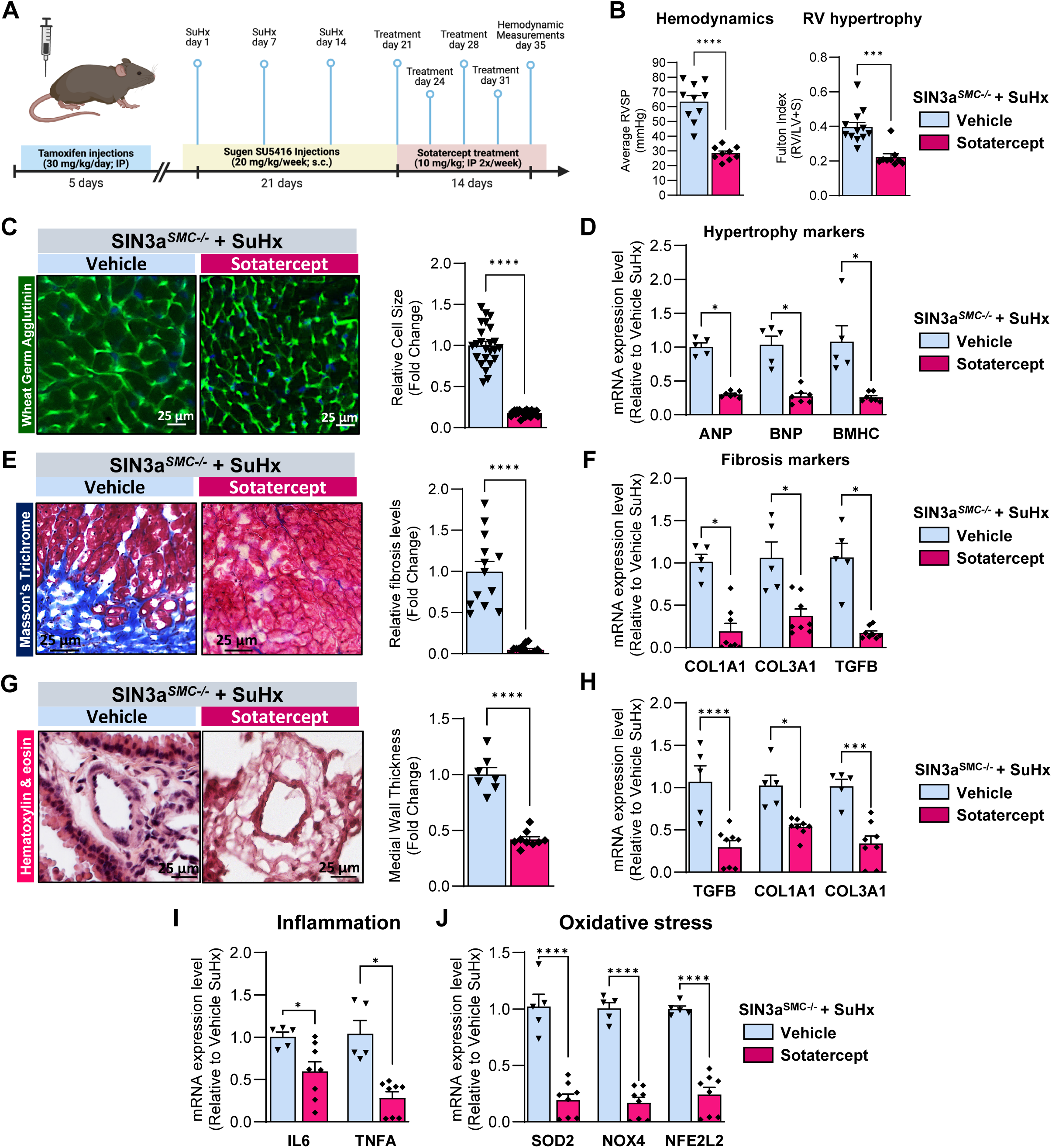
Sotatercept attenuates PAH in SIN3a^SMC-/-^ mice by restoring BMPR2/TGF-β signaling balance and suppressing HIF-1α. **(A)** Schematic overview of the experimental design used to evaluate the therapeutic efficacy of Sotatercept in the SuHx-induced PAH conditional SIN3a*^SMC-/-^* KO mouse model. Tissues were harvested 35 days after initiation of treatment for molecular and histological analyses. (**B**) RVSP (left panel) and RV hypertrophy assessed by the Fulton Index (RV/[LV+S]; right panel) in vehicle- and Sotatercept-treated SIN3a^SMC^ KO mice. Representative right ventricular sections stained with fluorescent WGA to assess cardiomyocyte cross-sectional area, with corresponding quantitative analysis shown to the right. qRT-PCR analysis of hypertrophic marker gene expression (*ANP, BNP, BMHC*) in RV tissue from the indicated groups. (**E**) Representative Masson’s trichrome-stained RV sections from vehicle- and Sotatercept-treated SIN3a*^SMC-/-^* KO mice. (**F**). qRT-PCR analysis of profibrotic gene expression (*TGFB, COL1A1, COL3A1*) in RV tissue from the indicated groups. (**G**). Representative hematoxylin and eosin (H&E)-stained lung sections from the indicated mice, with quantification of pulmonary arterial medial wall thickness shown on the right from vehicle- and Sotatercept-treated SIN3a*^SMC-/-^* KO mice. (**H**). qRT-PCR analysis of profibrotic marker expression in lung tissue from vehicle- and Sotatercept-treated SIN3a*^SMC-/-^* KO mice. (**I-J**). RT-PCR analysis of pro-inflammatory cytokines (*IL6, TNFA*) and oxidative stress–related genes (*NOX4, SOD2, NFE2L2*) in the lungs of Sotatercept-treated SIN3a*^SMC-/-^*KO mice compared with vehicle-treated controls. Data are presented as mean ±SEM; ns: not significant, * p<0.05, *** p < 0.001.

We next analyzed the expression of ID isoforms, and we observed that Sotatercept restored ID1, ID2, and ID3 mRNA levels in SIN3a*^SMC-/-^* KO-treated mice compared to vehicle-treated mice (**Figure 7A**). Sotatercept also reduced HIF1a and restored BMPR2 mRNA levels in SIN3a*^SMC-/-^*KO-treated mice compared to vehicle-treated mice (**Figure 7B**). Surprisingly, Sotatercept treatment restored SIN3a protein levels and significantly increased BMPR2 and p-SMAD1/5/9 levels (**Figure 7C-D**). Notably, Sotatercept treatment also reduced HIF1a expression level and TGFβ downstream SMAD2/3 activation in SuHx-exposed SIN3a*^SMC-/-^* KO mice (**Figure 7C-D**), highlighting a novel mechanism by which Sotatercept mitigates hypoxia-driven signaling in PAH. Our findings validate the role of SIN3a in regulating BMPR2 expression and downstream signaling, and provide, for the first time, compelling evidence that SIN3a critically influences profibrotic and pro-proliferative TGFB expression in PAH through HIF1a. Furthermore, Sotatercept treatment attenuated the effects of SIN3a loss in PASMCs and restored the balance between BMPR2/TGFB signaling (**Figure 7E**). Collectively, these results strongly support that SIN3a loss in PAH can be mitigated by Sotatercept, highlighting SIN3a as a clinically relevant therapeutic target for PAH. Overall, the loss of SIN3a in SMCs triggers a pathological cascade marked by increased inflammation, ECM remodeling, and oxidative stress by impairing the TGF-β/BMPR2 signaling.

**Figure 7.**
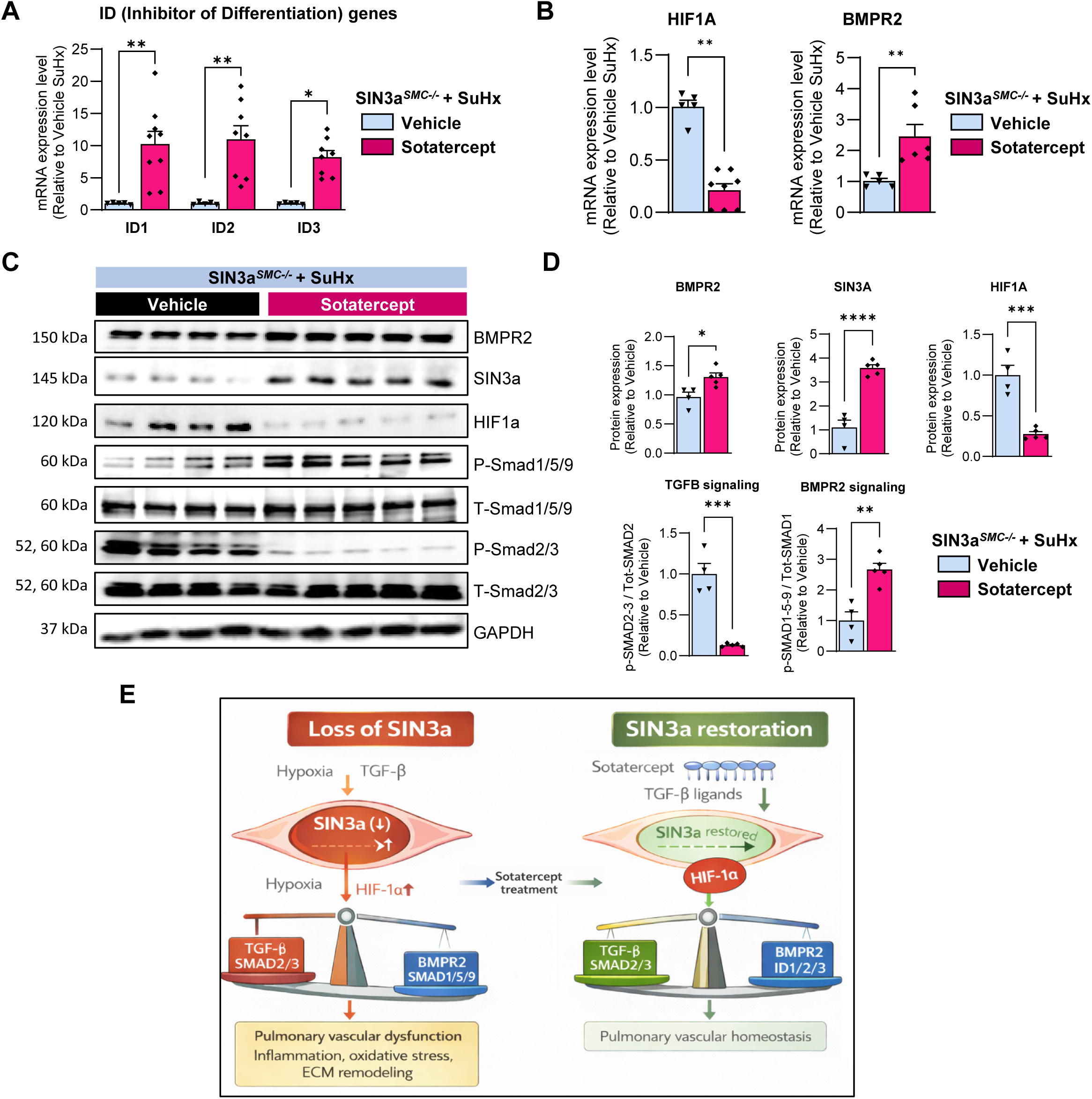
Restoration of BMPR2 and Suppression of TGF-β/HIF-1α Signaling by Sotatercept in SIN3a*^SMC-/-^* KO mice. (**A**) qRT-PCR analysis of *ID1, ID2*, and *ID3* mRNA expression in lungs from SIN3a*^SMC-/-^* KO lungs treated with Sotatercept compared to vehicle-treated controls. (**B**) qRT-PCR analysis of *HIF1α* and *BMPR2* mRNA expression in lungs from SIN3a*^SMC-/-^* KO lungs treated with Sotatercept compared to vehicle-treated controls. (**C-D**) Representative immunoblot analysis (**C**) and corresponding densitometric quantification (**D**) of SIN3a, BMPR2, SMAD signaling components, and HIF-1α protein levels in lungs from vehicle- and Sotatercept- treated SIN3a*^SMC-/-^* KO mice. (**E**) Schematic model illustrating the proposed mechanism by which SIN3a loss of function promotes PAH features and how Sotatercept treatment counteracts SIN3a deficiency. Sotatercept represses TGF-β signaling, HIF-1α expression, pro-inflammatory and oxidative stress pathways, while restoring BMPR2 signaling and pathway balance. Restoration of BMPR2/TGF-β signaling by SIN3a and Sotatercept limits oxidative stress, inflammation, and extracellular matrix remodeling through inhibition of HIF-1α–dependent pathways. Schematic Created with BioRender.com.

## DISCUSSION

PAH is increasingly recognized as a disease driven not only by genetic susceptibility and environmental stressors but also by maladaptive epigenetic reprogramming that stabilizes pathogenic cellular states. In this context, our findings identified SIN3a as a central epigenetic integrator that couple’s hypoxia sensing, growth factor signaling, and chromatin regulation to maintain pulmonary vascular homeostasis. Loss of SIN3a, observed in several hyperproliferative malignancies, emerges here as a mechanistic driver of pulmonary vascular remodeling, thereby extending cancer-associated epigenetic paradigms to PAH pathobiology.

SIN3a functions as a scaffold for multiprotein chromatin-modifying complexes by coordinating histone deacetylation, DNA methylation, and transcription factor accessibility. Its reduction in approximately 61% of non-small cell lung cancers ^41^ and its role in breast cancer progression via epigenetically mediated deregulation of growth-related genes ^42^ underscore its importance in constraining abnormal cellular proliferation. PAH shares multiple features with malignancy, including sustained PASMC proliferation, apoptosis resistance, metabolic reprogramming, and loss of tumor suppressor-like signaling ^43^. Our data validated and supported the concept that SIN3a deficiency represents a shared epigenetic vulnerability that enables these pathological phenotypes across distinct disease contexts.

A key advancement of this study is the demonstration that SIN3a expression is dynamically regulated by the pulmonary microenvironment. We showed that hypoxia and TGF-β1 signaling converge on HIF-1α to suppress SIN3a expression in PASMCs, providing a direct mechanistic link between environmental stress and epigenetic dysregulation. This observation aligns with the reduced SIN3a levels detected in PAH patient tissues and experimental models ^8,16,18^ and suggests that SIN3a loss is not merely a downstream consequence of disease but an active participant in disease initiation and progression. Importantly, this regulation appeared reversible, highlighting the therapeutic relevance of targeting epigenetic plasticity by targeting SIN3a.

The restoration of SIN3a reactivated a coordinated network of protective transcriptional programs that collectively restrained the pathological behavior of PASMCs. These include a feedback loop that suppresses HIF-1α accumulation, attenuates oxidative stress and inflammatory signaling, and inhibits pro-fibrotic gene expression. The convergence of these effects suggests that SIN3a acts not as a single-pathway regulator but as a higher-order transcriptional organizer that stabilizes nonpathological cellular identity. Therefore, loss of SIN3a appears to enable PASMCs to transition into a maladaptive state characterized by heightened stress responsiveness, inflammation, and fibrosis. This organizing role is consistent with emerging concepts in epigenetic regulation, in which chromatin-modifying complexes act as “state keepers” that preserve cellular identity under stress and prevent pathological phenotypic reprogramming. In this context, SIN3a deficiency may represent a critical epigenetic vulnerability that allows sustained environmental or metabolic stress to drive PASMC dysfunction and disease progression.

Interestingly, central to this protective function is SIN3a’s regulation of the BMPR2/TGF-β signaling balance, a critical determinant of pulmonary vascular remodeling. BMPR2-mediated SMAD1/5/9 signaling exerts antiproliferative and anti-fibrotic effects in the pulmonary vasculature, whereas excessive TGF-β signaling promotes PASMC hyperproliferation, ECM deposition, and vascular stiffening ^44^. Disruption of this balance, particularly in the setting of BMPR2 haploinsufficiency or loss-of-function mutations, is a defining feature of PAH. Our previous study demonstrated that SIN3a preserves BMPR2 expression by preventing promoter DNA and histone methylation ^8^. Here, we extend these findings by showing that SIN3a overexpression restores BMPR2 signaling and SMAD1/5/9 phosphorylation and ID isoforms expression through suppression of HIF-1α and TGF-β1 activity. Conversely, SIN3a loss shifts the signaling equilibrium toward unchecked TGF-β dominance, reinforcing pathological remodeling.

The interaction between hypoxia, HIF-1α, and TGF-β signaling further amplifies this pathogenic loop. HIF-1α is a master regulator of cellular adaptation to low oxygen tension and is stabilized under hypoxic conditions, which is typical of the remodeled pulmonary vasculature. HIF-1α activation enhances TGF-β/SMAD2/3 signaling and collagen deposition, which are attenuated by HIF-1α inhibition ^45^. Conversely, TGF-β has been shown to increase HIF-1α expression and transcriptional activity, establishing a feed-forward circuit that sustains pathological signaling^46^. Our data suggest that SIN3a is a critical epigenetic check on this circuit. Loss of SIN3a removes a transcriptional constraint on HIF-1α-driven programs, thereby locking PASMCs into a hypoxia-adapted and pro-remodeling state.

Our SIN3a*^SMC-/-^* KO model underscores the in vivo relevance of this mechanism. When subjected to SuHX stress, these mice exhibited exaggerated pulmonary vascular and RV remodeling, demonstrating that SIN3a loss is sufficient to sensitize the vasculature to environmental and inflammatory stressors. This finding supports a two-hit model in which epigenetic vulnerability conferred by SIN3a deficiency amplifies the pathogenic impact of hypoxia and growth factor imbalance, thereby accelerating disease progression.

From a therapeutic perspective, our findings provide important new insights into the disease-modifying actions of Sotatercept. Sotatercept, an activin receptor type IIA–Fc fusion protein, restores BMP signaling by sequestering pathogenic TGF-β superfamily ligands and has demonstrated robust clinical efficacy in PAH ^47^. Although its ability to rebalance BMP and TGF-β signaling is well established, we deliberately treated SIN3a*^SMC-/-^* KO mice with Sotatercept as a monotherapy to isolate its direct effects on PAH pathology. Strikingly, Sotatercept not only attenuated pulmonary vascular and RV remodeling but also restored SIN3a expression in SIN3a*^SMC-/-^*KO mice, an effect likely mediated by suppression of HIF-1α activity and consequent inhibition of pathogenic TGF-β signaling. This previously unrecognized epigenetic action suggests that Sotatercept extends beyond transient pathway modulation and instead promotes reprogramming of maladaptive transcriptional states toward a more stable, homeostatic phenotype. Such epigenetic normalization offers a compelling mechanistic explanation for the durability of Sotatercept’s clinical benefits and positions restoration of epigenetic control as a key therapeutic objective in PAH. Collectively, these findings redefine PAH as a disease sustained by epigenetic dysregulation at the intersection of hypoxia signaling and growth factor imbalance. SIN3a has emerged as a central node that integrates environmental stress with chromatin-based transcriptional control, and its loss represents a convergence point for multiple pathogenic pathways. By identifying SIN3a as both a mechanistic driver of disease and a downstream effector of therapeutic intervention, this study opens new avenues for precision medicine strategies aimed at restoring epigenetic stability and halting pulmonary vascular remodeling.

### Conclusion

Our findings establish SIN3a as a central epigenetic regulator of pulmonary vascular homeostasis, and its deficiency is sufficient to exacerbate PAH. Hypoxia and TGF-β-driven HIF-1α activation suppress SIN3a expression, disrupt protective transcriptional programs, imbalance BMPR2/TGF-β signaling, and amplify oxidative stress, inflammation, fibrosis, and PASMC overgrowth. Restoration of SIN3a reverses these maladaptive responses, underscoring its critical role in constraining pathological vascular remodeling. Importantly, we identified the restoration of SIN3a as a previously unrecognized component of Sotatercept’s disease-modifying effects. By suppressing HIF-1α activity and re-establishing SIN3a-dependent epigenetic control, Sotatercept not only rebalances BMP and TGF-β signaling but also targets a fundamental driver of pulmonary vascular remodeling. Together, these findings establish the SIN3a-HIF-1α-BMPR2/TGF-β axis as a central pathogenic hub in PAH and identify epigenetic modulation of this pathway as a rational and promising strategy for precision therapies aimed at halting and potentially reversing disease progression.

## ABREVIATION LIST

α-SMA: Alpha-smooth muscle actin
ANP: Atrial natriuretic peptide
BMPR2: Bone morphogenetic protein receptor type 2
BNP: Brain natriuretic peptide
COL1A1: Collagen type I alpha 1
COL3A1: Collagen type III alpha 1
CreER^T2^: Tamoxifen-inducible Cre recombinase
CTGF: Connective tissue growth factor
DEG: Differentially expressed gene
ECM: Extracellular matrix
EMT: Epithelial-to-mesenchymal transition
GAGE: Generally Applicable Gene-set Enrichment
GAPDH: Glyceraldehyde 3-phosphate dehydrogenase
GO: Gene Ontology
H&E: Hematoxylin and eosin
HIF1α: Hypoxia-inducible factor 1-alpha
IL6: Interleukin 6
KO: Knockout
LV+S: Left ventricle plus septum
MCT: Monocrotaline
NF-κB: Nuclear factor kappa-light-chain-enhancer of activated B cells
NFE2L2: Nuclear factor erythroid 2–related factor 2
NOX4: NADPH oxidase 4
PAH: Pulmonary arterial hypertension
PASMC: Pulmonary artery smooth muscle cell
PH: Pulmonary hypertension
ROS: Reactive oxygen species
RV: Right ventricle
RVSP: Right ventricular systolic pressure
SIN3a: Switch-independent 3A
SMAD1/5/9: Mothers against decapentaplegic homolog 1/5/9
SMAD2/3: Mothers against decapentaplegic homolog 2/3
SMC: Smooth muscle cell
SMMHC: Smooth muscle myosin heavy chain
SuHx: Sugen 5416 plus hypoxia
TGF-β: Transforming growth factor-beta
TGFBR1: TGF-β receptor 1
TNF-α: Tumor necrosis factor-alpha
VEGFR2: Vascular endothelial growth factor receptor 2
WGA: Wheat germ agglutinin

## Availability of data and materials

The datasets generated and analyzed during the current study are available from the corresponding author upon reasonable request. RNA sequencing data have been deposited in a publicly accessible repository prior to publication.

## Ethics approval and consent to participate

All animal experiments were approved by the Institutional Animal Care and Use Committee (IACUC) of the Icahn School of Medicine at Mount Sinai and were conducted in accordance with the NIH Guide for the Care and Use of Laboratory Animals.

## Consent for publication

This study was exempt from ethical review as it did not involve individual data requiring consent.

## Acknowledgments

Not applicable. All the contributions to this article are limited to the authors listed in the manuscript.

## Authors contributions

K.J., A.G., M.T.O., and S.Z. performed the experiments and collected and analyzed the data. G.D. contributed to the study design and provided the essential experimental resources. I.T. contributed to manuscript editing and critical revision of intellectual content. M.B. and L.H. conceived and designed the study, supervised the research, performed the experiments, analyzed and interpreted the data, and wrote the manuscript. All authors have reviewed and approved the final manuscript.

## Supplementary Figures

**Supplementary Figure 1. Global RNA-sequencing quality control and differential expression analysis.** (**A**) Principal component analysis (PCA) of RNA-sequencing data demonstrates clear separation between Ad-SIN3a- and Ad-Control-infected PASMCs, indicating a global shift in the transcriptomic landscape following SIN3a overexpression. (**B**) MA plot showing differentially expressed genes (DEGs) between SIN3a-overexpressing and control PASMCs. Significantly upregulated and downregulated transcripts are highlighted.

**Supplementary Figure 2. Gene Ontology enrichment of SIN3a-regulated transcriptional clusters.** (**A**) Gene Ontology (GO) enrichment analysis of SIN3a-regulated transcripts identifies four major biological clusters, including cellular stress responses, RNA processing, signaling pathways, and ECM organization. (**B**) WikiPathways enrichment analysis reveals diverse regulatory modules altered by SIN3a overexpression, including nuclear receptor signaling and apoptotic pathways, highlighting the multifaceted transcriptional role of SIN3a.

**Supplementary Figure 3. Network-level integration of SIN3a-regulated transcriptional programs.** Network topology map illustrating functional interactions among SIN3a-regulated biological processes. Upregulated pathways are shown in green, and downregulated pathways are shown in red. The network reveals extensive crosstalk among ECM remodeling, transcriptional regulation, RNA metabolism, and cellular stress response pathways, underscoring SIN3a’s role as an integrative regulator of PASMC homeostasis.

**Supplementary Figure 4. SIN3a transcriptional targets converge on pathogenic signaling modules implicated in PAH.** (**A**) GO enrichment analysis identifies key biological processes modulated by SIN3a, including oxidative stress responses, extracellular matrix organization, pro-inflammatory signaling, cell migration and adhesion, and ECM remodeling-processes central to vascular pathology. **(B)** GO enrichment analysis of the top 250 SIN3a-downregulated DEGs reveals unresolved cellular stress pathways, including the unfolded protein response (UPR), IL-2/IL-6 cytokine signaling, and chronic hypoxic injury responses.

**Supplementary Figure 5. Hallmark pathway enrichment of SIN3a-regulated genes.**

**(A)** Hallmark pathway enrichment analysis of the top 250 genes upregulated by SIN3a shows enrichment of epithelial-mesenchymal transition (EMT), hypoxia signaling, glycolysis, cholesterol biosynthesis, apoptosis, and inflammatory signaling pathways. **(B)** Hallmark pathway enrichment analysis of the top 250 genes downregulated by SIN3a demonstrates suppression of unresolved cellular stress pathways, including UPR, IL-2/IL-6 cytokine signaling, and chronic hypoxic injury responses.

**Supplementary Figure 6. Pro-fibrotic and pro-inflammatory signaling pathways regulated by SIN3a.** GSEA enrichment plots demonstrate suppression of ECM-related processes, including extracellular structure organization and collagen biosynthesis, in SIN3a-overexpressing PASMCs. In addition, genes regulated by SIN3a are enriched for ROS signaling, glycolysis, RNA splicing, EMT, hypoxia signaling, DNA repair, and stress-adaptive pathways, suggesting coordinated transcriptional and post-transcriptional regulation.

**Supplementary Figure 7. HIF-1α binding at promoter regions of SMAD3 and SMAD4.** ENCODE ChIP-seq analysis reveals HIF-1α binding at the promoter regions of SMAD3 and SMAD4.

**Supplementary Figure 8. SIN3a modulates IL-6 expression under hypoxic and TGF-β1 conditions.** qRT-PCR analysis showing modulation of IL-6 mRNA expression in FD-PASMCs cultured under normoxic conditions or exposed to hypoxia (1% O₂) and TGF-β1, with or without SIN3a overexpression, n=3. Data are presented as mean ±SEM; *** p < 0.001.

**Supplementary Figure 9. Specific conditional deletion of SIN3a in smooth muscle cells. (A)** SIN3a^flox/flox^ mice were crossed with SMMHC-Cre⁺ mice to enable tamoxifen-inducible, SMC-specific deletion of SIN3a. Adult male mice (8-10 weeks old) received daily intraperitoneal injections of tamoxifen (35 mg/kg/day; Sigma-Aldrich) dissolved in corn oil for five consecutive days. DNA genotyping confirmed SIN3a deletion in SIN3a^SMC-/-^ mice compared with floxed littermates. Only male mice were used to ensure uniform Cre expression and recombination efficiency, as the Myh11-Cre/ERT2 transgene is located on the Y chromosome. Littermates lacking the Cre allele or not receiving tamoxifen served as controls. **(B)** qRT-PCR analysis shows no significant change in SIN3b mRNA expression in SIN3a*^SMC-/-^*mice compared with littermate controls. Data are presented as mean ±SEM; ns: not significant.

**Supplementary Figure 10. IL6 and IL-1β mRNA expression in SIN3a-deficient mice.** qRT-PCR analysis of IL-1β mRNA expression levels in SIN3a*^SMC-/-^* and littermate control mice under normoxic and Sugen–hypoxia (SuHx) conditions, n=6. Data are presented as mean ±SEM; ns: not significant.

**Supplementary Figure 11. Mesenchymal and contractile marker expression in SIN3a-deficient mice.** qRT-PCR analysis of α-SMA, SLUG, and SNAI1 mRNA expression levels in SIN3a*^SMC-/-^* and littermate control mice under normoxic and SuHx conditions, n=6. Data are presented as mean ±SEM; ns: not significant, * p<0.05, **p<0.01, *** p < 0.001.

