## Supplementary figures and images for "Sotatercept Reverses SIN3a Deficiency-Driven PAH by Reprogramming BMPR2/TGF-β-HIF-1α Signaling Pathways"

### Supplementary Figure 1

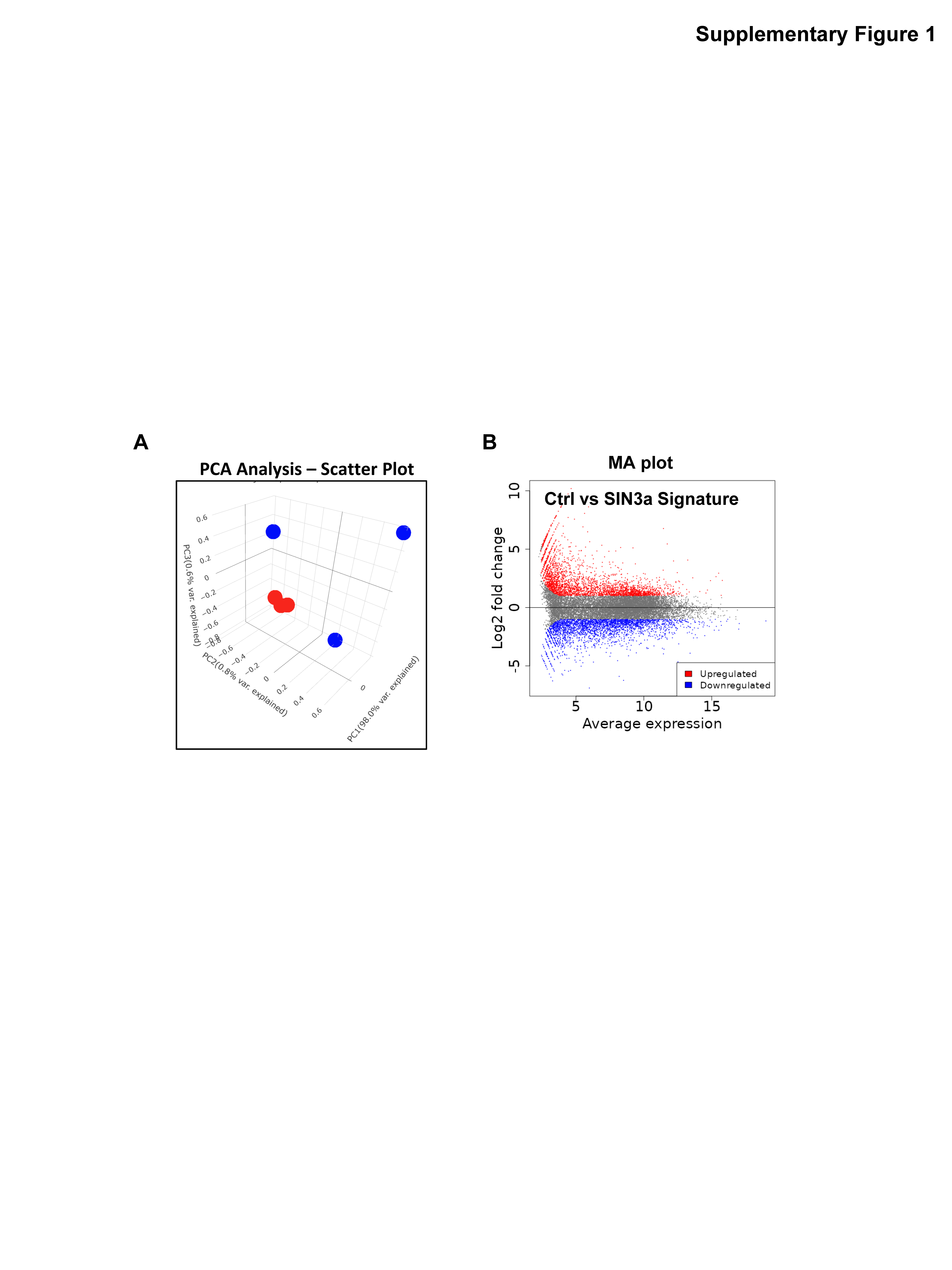

### Supplementary Figure 2

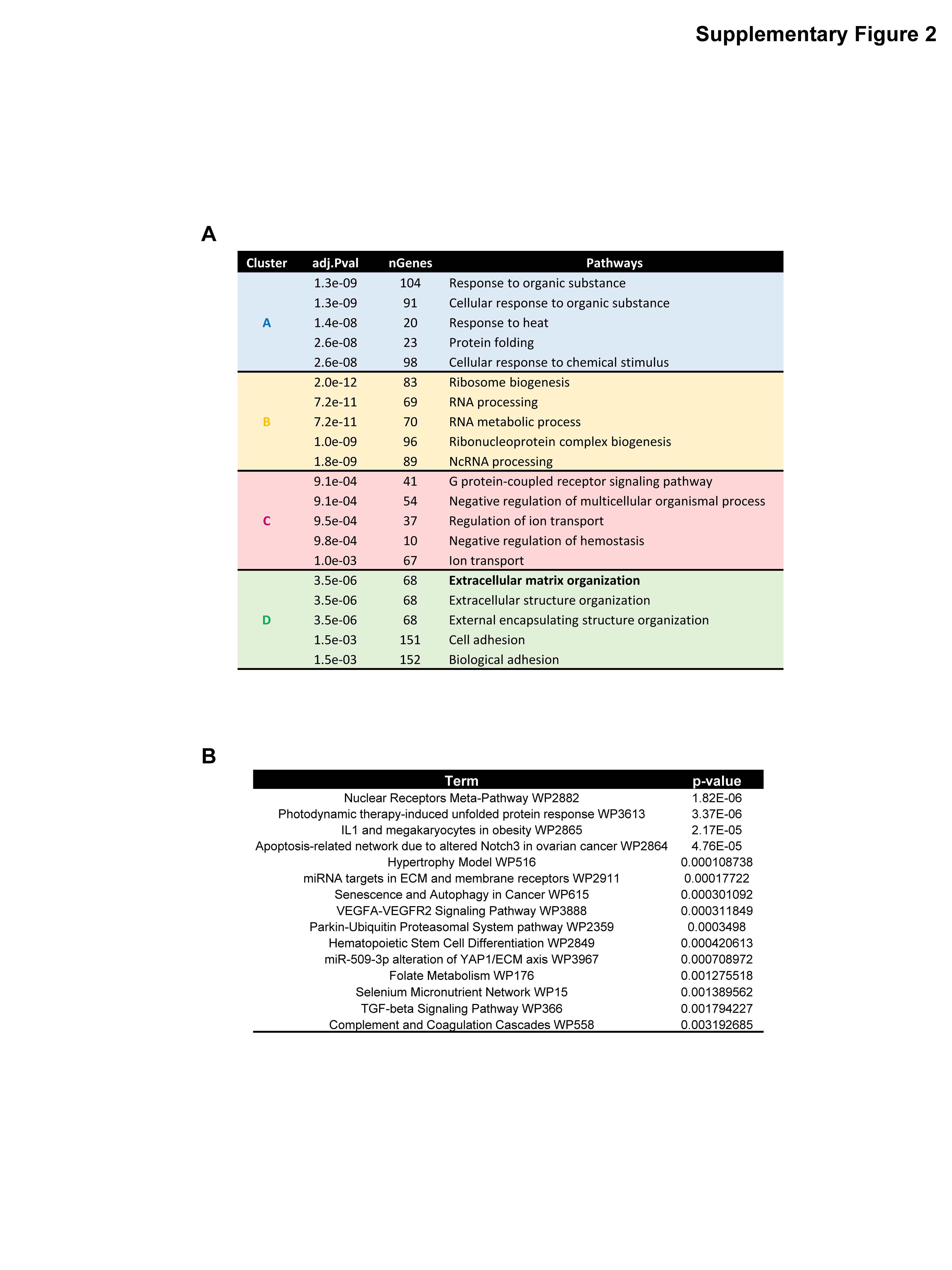

### Supplementary Figure 3

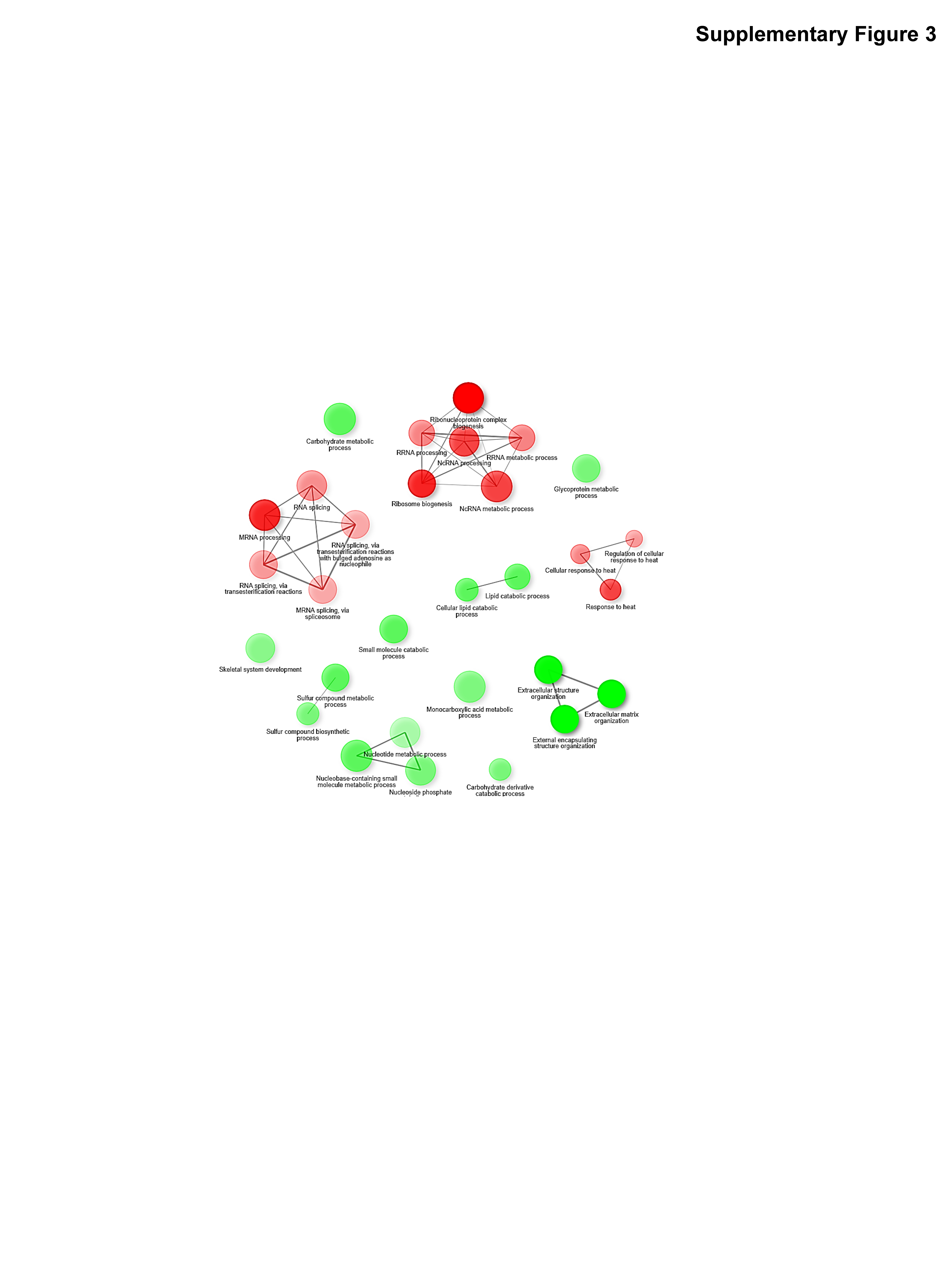

### Supplementary Figure 4

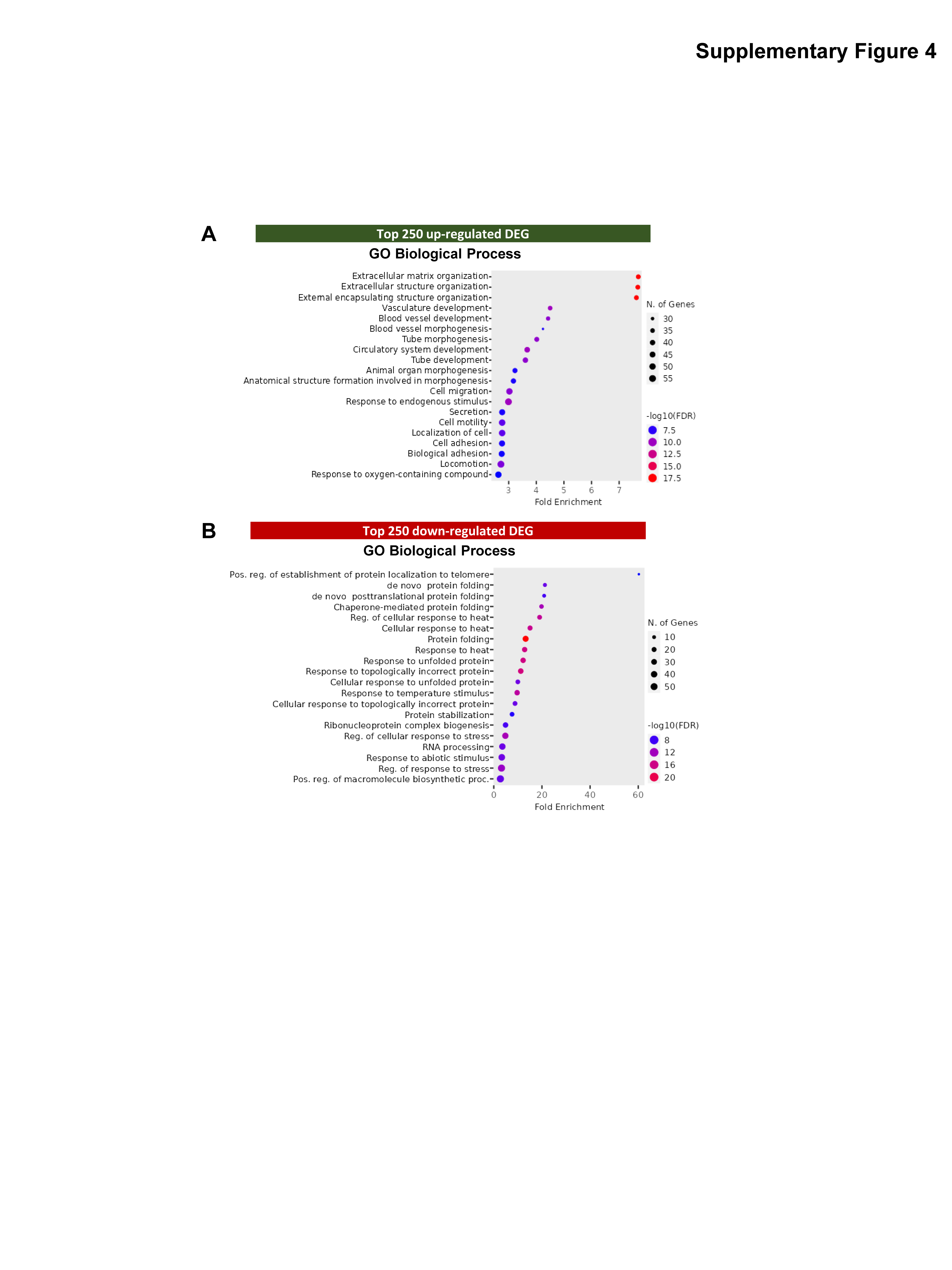

### Supplementary Figure 5

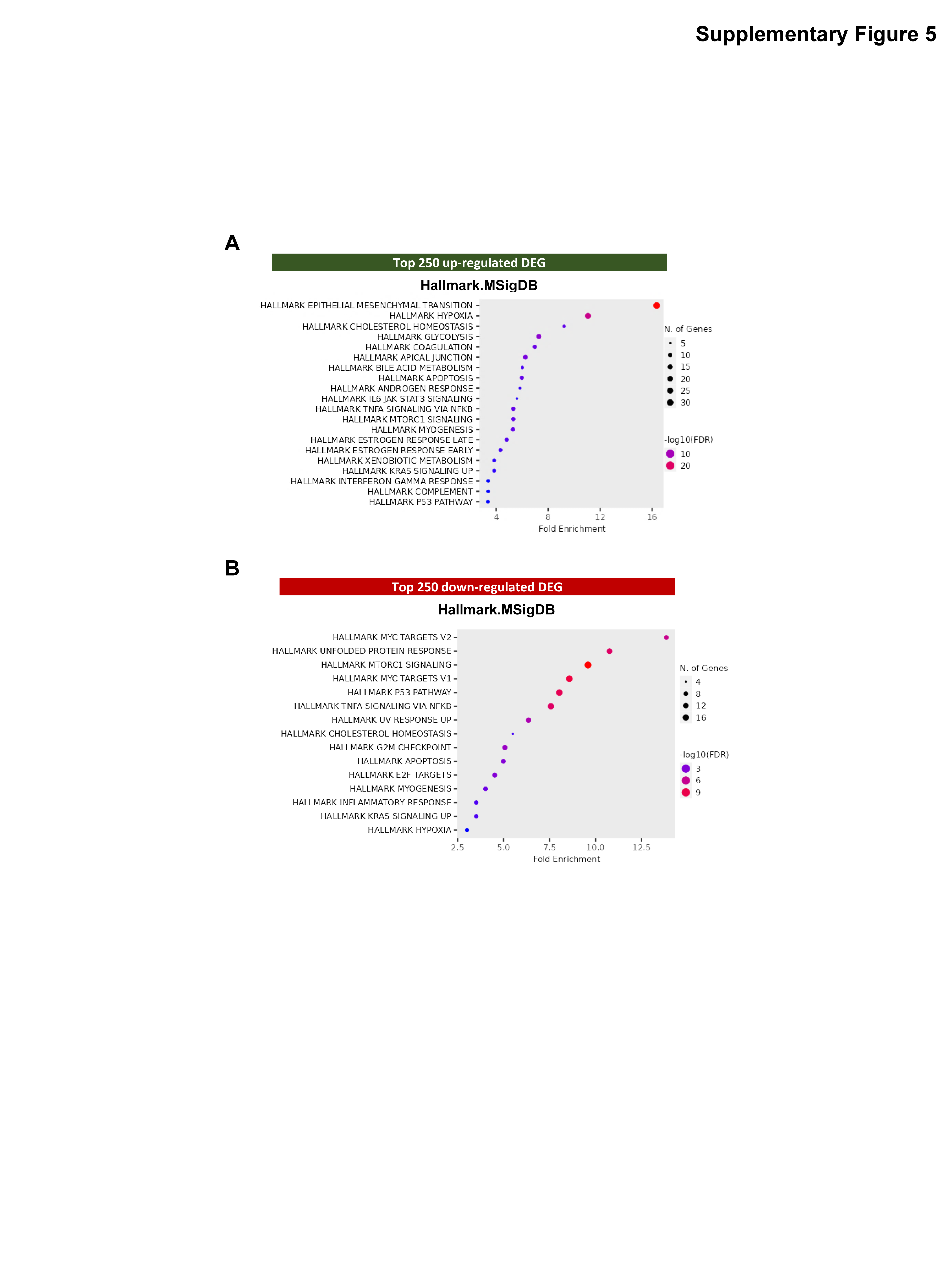

### Supplementary Figure 6

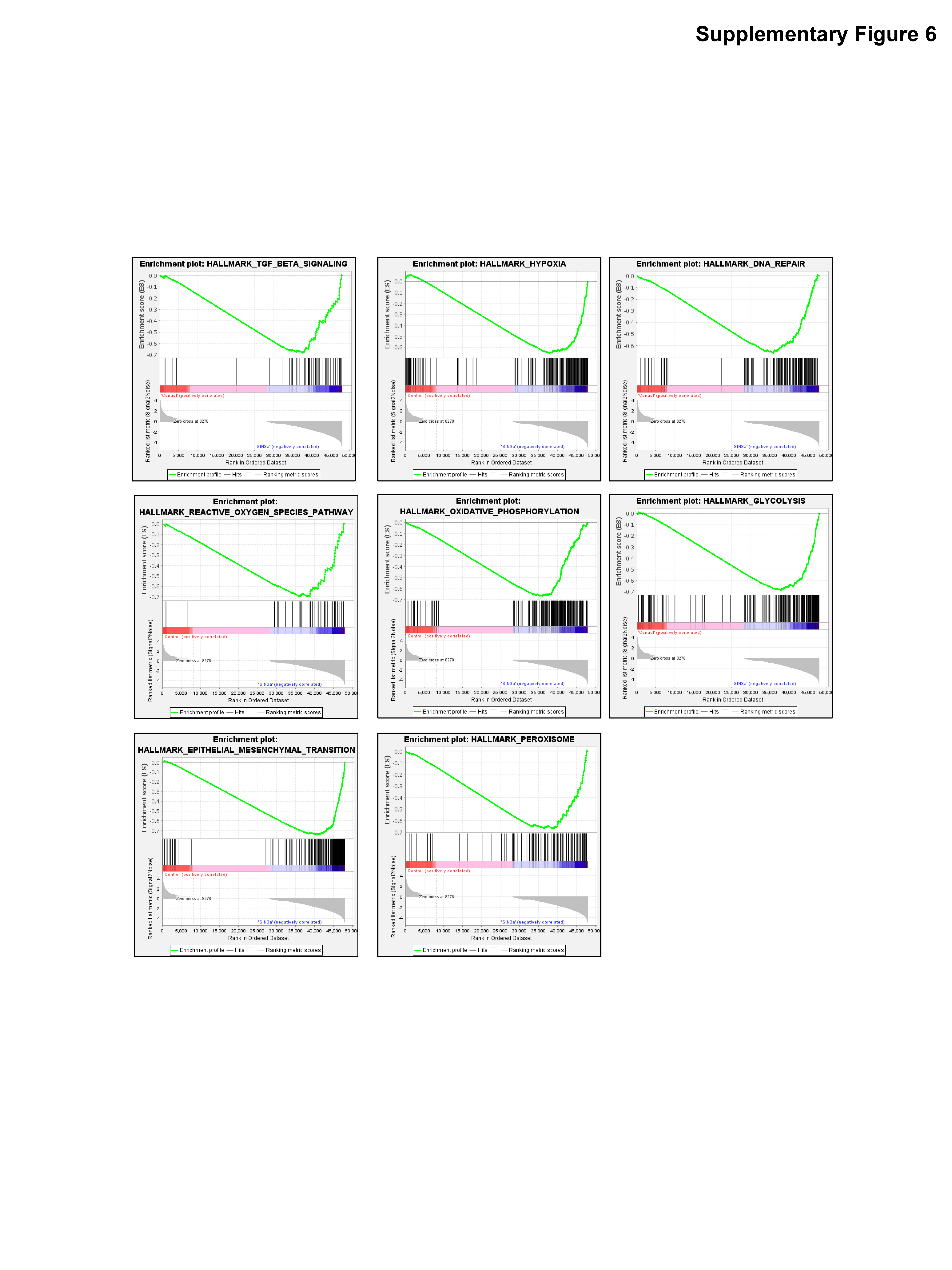

### Supplementary Figure 7

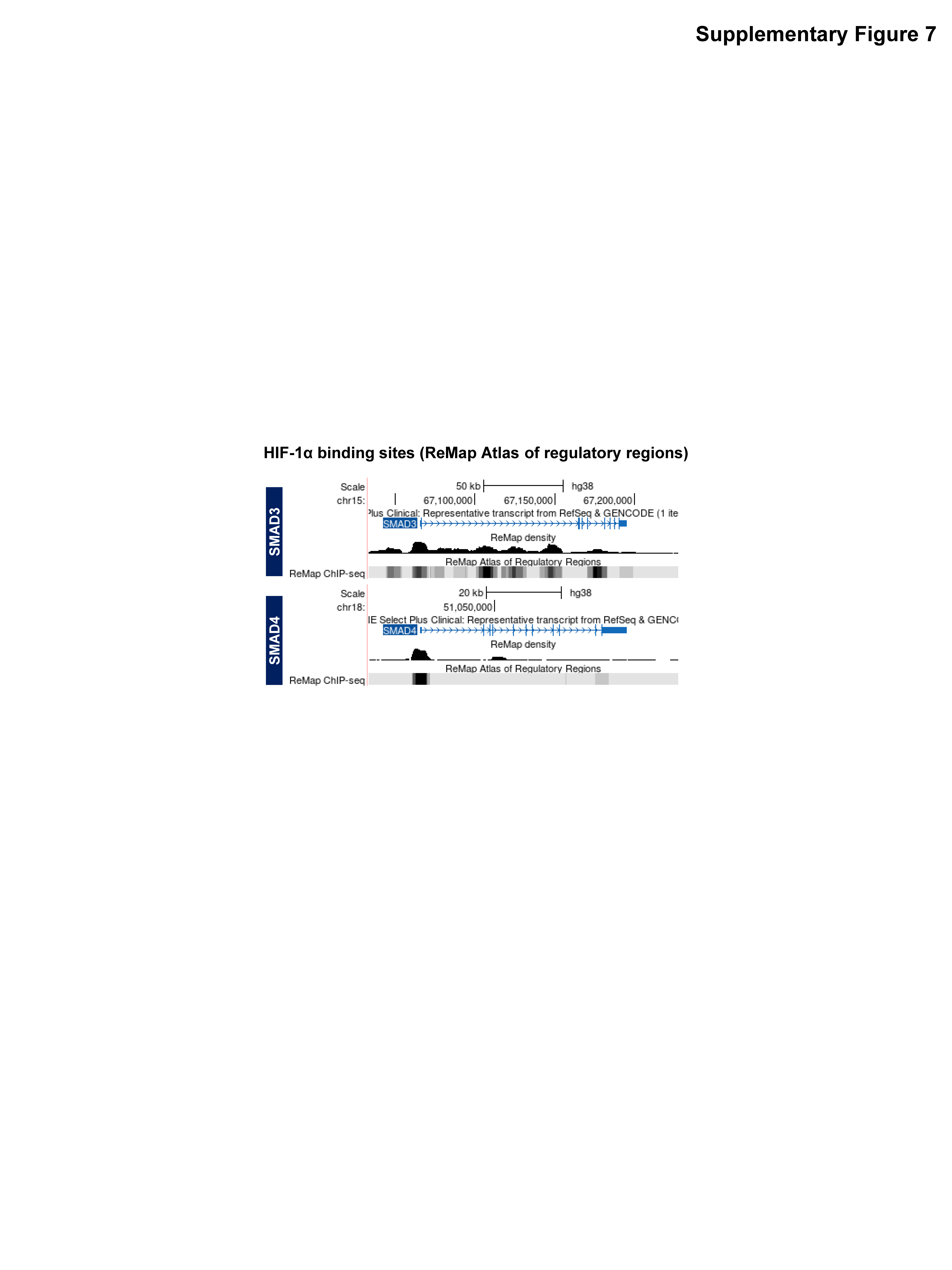

### Supplementary Figure 8

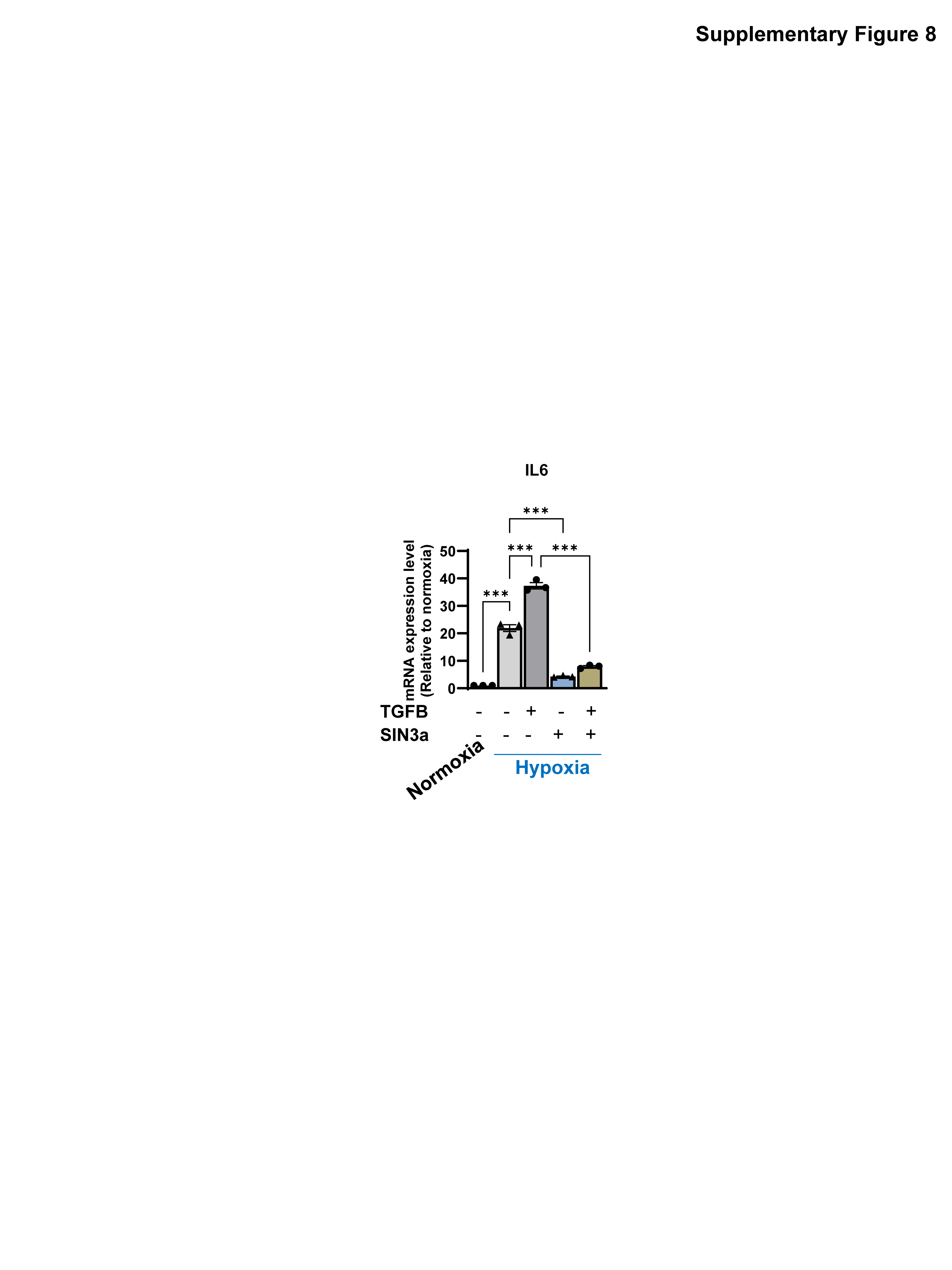

### Supplementary Figure 9

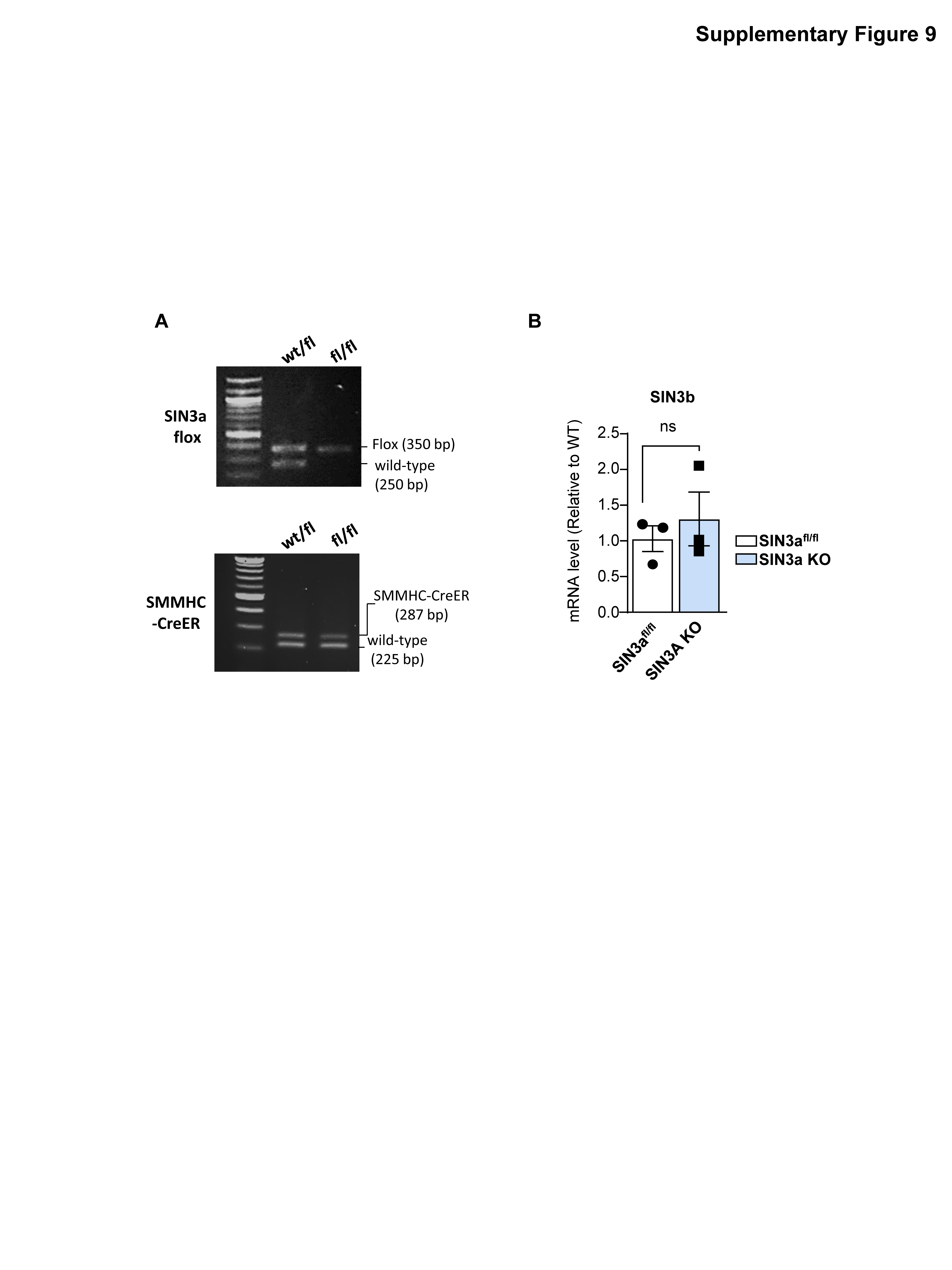

### Supplementary Figure 10

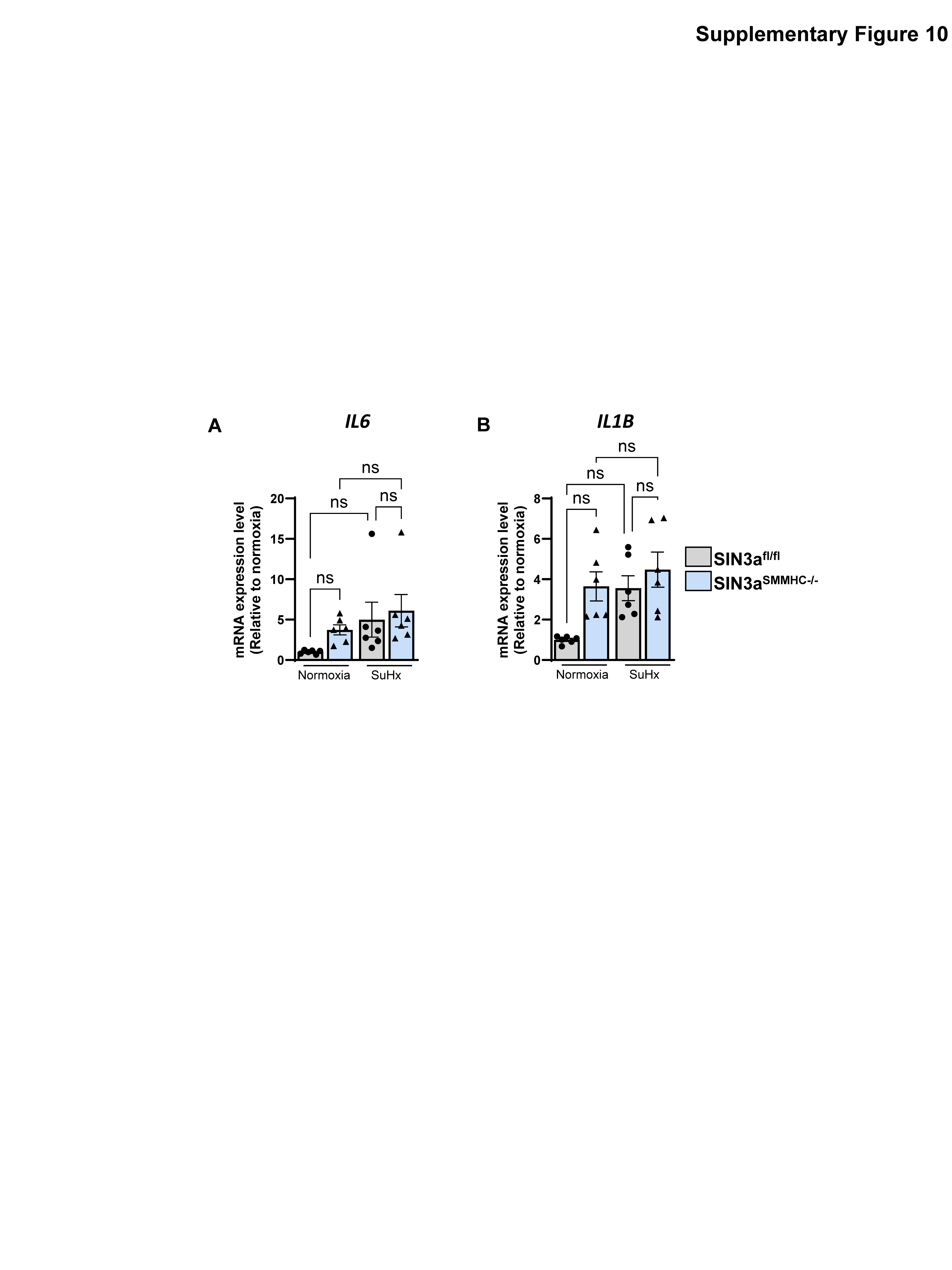

### Supplementary Figure 11

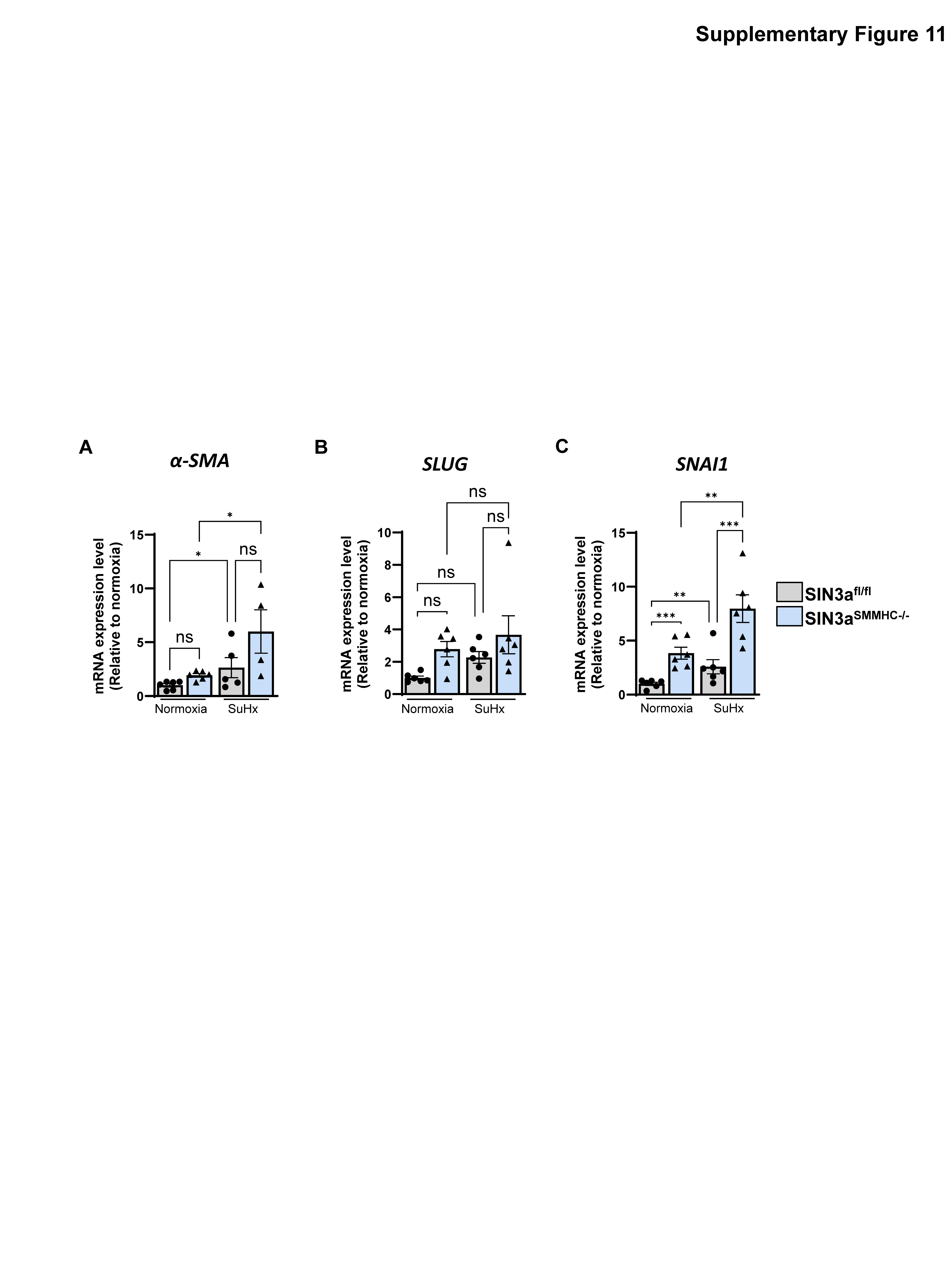
