## Supplementary Table 1 for "Sotatercept Reverses SIN3a Deficiency-Driven PAH by Reprogramming BMPR2/TGF-β-HIF-1α Signaling Pathways"

**Supplementary Table 1.** Primer sequences for RT-qPCR analysis, antibodies used, source, and concentration of pharmacological treatments used.

| **Application** | **Gene symbol** | **Species** | **Forward primer (5′-3′)** | **Reverse primer (5′-3′)** |
| --- | --- | --- | --- | --- |
| **RT-qPCR** | SIN3A | Human | TCAACCACCACCCCAACATC | CTGGCCGGAGTATGTGCTTG |
|  | SIN3A | Mouse | GATTCCAGGGCCAACCAGAC | AGCCACCTGGGCATACACCT |
|  | SIN3B | Mouse | GTTGTCACGGATGGCTCCTG | CTCCTCCTCCTTGGCCTTCA |
|  | BMPR2 | Human  Mouse | TTTACAGAGTGCCTTTGATGGAAC | ATTCTGTGTGAAGATAAGCCAGTC |
|  | HIF1A | Human  Mouse | GCT TGG TGC TGA TTT GTG AAC C | CCT GTG GTG ACT TGT CCT TTA G |
|  | COL1A1 | Human  Mouse | TTCAGTGGTTTGGATGGTGCCAA | CCAGCTTCACCCTTAGCACCA |
|  | COL3A1 | Human  Mouse | GAGATGTCTGGAAGCCAGAACCATG | ATCTCCCTTGGGGCCTTGAGGT |
|  | TGFB | Human  Mouse | CCTGCAAGACCATCGACATGGAG | GGTCGCGGGTGCTGTTGTA |
|  | ANP | Mouse | GCTTCCAGGCCATATTGGAG | GGGGGCATGACCTCATCTT |
|  | BNP | Mouse | CTGGGAAGTCCTAGCCAGTC | TTTTCTCTTATCAGCTCCAGCA |
|  | β-MHC | Mouse | ACTGTCAACACTAAGAGGGTCA | TTGGATGATTTGATCTTCCAGGG |
|  | SOD2 | Mouse | TGGAGAACCCAAAGGAGAGTTG | AGCGACCTTGCTCCTTATTGAA |
|  | NOX4 | Human  Mouse | GTGAACATCCAGCTGTACCTCA | GTCCACAGCAGAAAACTCCAAC |
|  | NFE2L2 | Human  Mouse | TGCCCCTGGAAGTGTCAAACA | GGCTTGAATGTTTGTCTTTTGTGAATGG |
|  | IL-6 | Human | ACAAGAGTAACATGTGTGAAAGCAG | ACTCTCAAATCTGTTCTGGAGGTAC |
|  |  | Mouse | GGTCTTCTGGAGTTCCGTTTCT | AGAGCATTGGAAGTTGGGGTAG |
|  | TNFA | Mouse | GGTGCCTATGTCTCAGCCTC | ACTGATGAGAGGGAGGCCAT |
|  | IL1B | Mouse | TGGACCTTCCAGGATGAGGACA | GTTCATCTCGGAGCCTGTAGTG |
|  | ID1 | Mouse | TGCTCTACGACATGAACGGCT | TCTCGCCGTTCAGGGTG |
|  | ID2 | Mouse | TTAGGAAAAACAGCCTGTCGGAC | CTTCTTGTTCTGGGGGATGCTG |
|  | ID3 | Mouse | TCAGCTTAGCCAGGTGGAAATC | TTGGAGATCACAAGTTCCGGAG |
|  | ASMA | Mouse | CATGAGCCGTGAAGTGCAGT | TGAGCCACCTGTTCCATCTG |
|  | SLUG | Mouse | CCCCATGCCATTGAAGCTGA | GCCTTGCCACAGATCTTGCA |
|  | SNAI1 | Mouse | GATGCACATCCGAAGCCACA | TGTGGAGCAAGGACATTCGG |
|  | 18S | Mouse  Human | GTAACCCGTTGAACCCCATT | CCATCCAATCGGTAGTAGCG |
| **Genotyping PCR** | Sin3A-1 | Mouse | GTC CTC AGG GAA GAC GTT GA |  |
|  | Sin3A-4 | Mouse | GCC CTG TCC TAT CTT GAC CA |  |
|  | Sin3A-5 | Mouse | AGG ACC ACC AAA GTT CAG GA |  |
|  | SMMHC-CreERT2-Wt | Mouse | TGACCCCATCTCTTCACTCC | AACTCCACGACCACCTCATC |
|  | SMMHC-CreERT2-Mutant | Mouse | AGTCCCTCACATCCTCAGGTT |  |
| **Immunoblotting** | **Protein symbol** | **Antibody source** | | **Dilution** |
|  | SIN3A | Cell signaling | | 1:1000 |
|  | HIF1A | Cell signaling | | 1:1000 |
|  | BMPR2 | Invitrogen | | 1:1000 |
|  | Phospho-SMAD1/5/9 | Cell signaling | | 1:1000 |
|  | Phospho-SMAD2/3 | Cell signaling | | 1:1000 |
|  | Total-SMAD1 | Cell signaling | | 1:1000 |
|  | Tota-SMAD2/3 | Cell signaling | | 1:1000 |
|  | Β-ACTIN | Cell signaling | | 1:1000 |
|  | GAPDH | Invitrogen | | 1:5000 |
| **Pharmacological agents** | **Compounds** | **Concentration** | | **Source** |
|  | TGF-beta | 5 nM | | Peprotech |
|  | SU5416 | 20 mg/Kg | | MedChem Express |
|  | Sotatercept | 5 nM | | MedChem Express |
